# Prenatal alcohol exposure disrupts Shh pathway and primary cilia genes in the mouse neural tube

**DOI:** 10.1101/649673

**Authors:** Karen E. Boschen, Eric W. Fish, Scott E. Parnell

## Abstract

Neurulation-stage alcohol exposure (NAE; embryonic day [E] 8-10) is associated with midline craniofacial and CNS defects that likely arise from disruption of morphogen pathways, such as Sonic hedgehog (Shh). Notably, midline anomalies are also a hallmark of genetic ciliopathies such as Joubert syndrome. We tested whether NAE alters Shh pathway signaling and the number and function of primary cilia, organelles critical for Shh pathway transduction. Female C57BL/6J mice were administered two doses of alcohol (2.9 g/kg/dose) or vehicle on E9. Embryos were collected 6, 12, or 24 hr later, and changes to Shh, cell cycle genes, and primary cilia were measured in the rostroventral neural tube (RVNT). Within the first 24 hours post-NAE, reductions in Shh pathway and cell cycle gene expression and the ratio of Gli3 forms in the full-length activator state were observed. RVNT volume and cell layer width were reduced at 12 hr. In addition, expression of multiple cilia-related genes were observed at 6 hr post-NAE. As a further test of cilia gene-ethanol interaction, mice heterozygous for *Kif3a* exhibited perturbed behavior during adolescence following NAE compared to vehicle-treated mice, and *Kif3a* heterozygosity exacerbated the hyperactive effects of NAE on exploratory activity. These data demonstrate that NAE downregulates the Shh pathway in a region of the neural tube that gives rise to alcohol-sensitive brain structures and identifies disruption of primary cilia function, or a “transient ciliopathy”, as a possible cellular mechanism of prenatal alcohol pathogenesis.

## Introduction

Prenatal alcohol exposure is the leading cause of preventable birth defects in the US, with Fetal Alcohol Spectrum Disorders (FASD) estimated to affect at least 5% of live births each year (1). Alcohol exposure often occurs before pregnancy is identified, during the early, important stages of embryonic development, such as gastrulation and neurulation. Early gestational alcohol exposure has been linked to growth retardation, central nervous system dysfunction, and distinct craniofacial malformations, including the “classic” facial phenotype of Fetal Alcohol Syndrome (FAS) characterized by midline defects such as hypotelorism and an absent philtrum (2, 3), as well as more subtle facial variances (4). Alcohol exposure during gastrulation, the stage when the embryo first forms distinct cell layers (3^rd^ week in humans, embryonic day [E] 7 in mice), induces widespread cell death in the neuroectoderm (5), and subsequent diminished Sonic hedgehog (Shh) signaling as a pathogenic mechanism for gastrulation-stage alcohol exposure (6–10). Alcohol exposure during neurulation, as the neural tube forms and closes (∼4^th^-5^th^ weeks in humans, E8-10 in mice), produces midline structural defects in brain regions such as the hypothalamus, ventricles, pituitary, and septal regions (11–14). However, while prenatal alcohol exposure during neurulation causes cell death in regions such as the rhombencephalon, alcohol-induced apoptosis is not as pronounced in the rostroventral neural tube (5), the portion of the neural tube that gives rise to ventral midline brain structures.

The Shh signaling pathway is transduced within immotile sensory organelles known as primary cilia (15, 16). Genetic disruption of the function and/or stability of primary cilia induces several developmental abnormalities, exemplified by the ciliopathy diseases, such as Joubert syndrome. Genetic ciliopathies cause multiple organ system defects, ocular malformations, cleft palates and lips, holoprosencephaly, and polydactyly (17, 18). Abnormal Shh signaling has been reported in cultured cells collected from humans with gene mutations linked to ciliopathies (19, 20) and knockdown of *Kif3a* or other intraflagellar transport proteins in mouse models of ciliopathies disrupts the Shh pathway (21–23) and causes Gli-dependent midfacial defects (24–26).

We hypothesized that neurulation-stage alcohol exposure (NAE) also disrupts Shh signaling in the rostroventral region of the neural tube (RVNT) that gives rise to many of the affected midline brain structures, explaining the manifestation of subtle craniofacial and CNS defects associated with alcohol exposure during this developmental window. We tested the effect of mid-neurulation (E9.0) alcohol exposure on expression of the *Shh* pathway and cell cycle genes within 24 hr following exposure. Following determination that NAE significantly decreased Shh and cell cycle gene expression, we were interested in whether neurulation-stage alcohol also affects primary cilia, as Shh transduction takes place within the primary cilia. To test this hypothesis, we analyzed the density of primary cilia in the RVNT and expression of genes known to play a role in cilia protein trafficking and ciliogenesis. Finally, we studied whether NAE interacts with cilia function to increase the sensitivity to the long-term effects of NAE. For this experiment, we exposed embryonic mice with a partial deletion of the key cilia gene *Kif3a*, which is a well-characterized mouse model of genetic ciliopathies, to E9.0 alcohol and then measured adolescent behavioral performance on tasks known to be affected by prenatal alcohol.

## Methods and Materials

### Animals

Male and female adult C57BL/6J mice were obtained from Jackson Laboratories (Stock No: 000664; Bar Harbor, ME). Males were housed singly and females were housed in groups of up to five per standard polycarbonate cage with cob bedding, a shelter, and nesting material. All mice had *ad libitum* access to food (Prolab Isopro RMH 3000, LabDiet, St. Louis, MO) and water, and were maintained on a 12:12 hr light/dark cycle. For mating, 1-2 female mice were placed into the cage of a male for 1-2 h and were checked for a vaginal plug to confirm copulation. E0.0 was defined as the beginning of the mating session when the plug was found. Mated females were weighed and housed in a clean cage with up to 5 mice. All experimental procedures and euthanasia protocols were approved by the Institutional Care and Use Committee (IACUC) at University of North Carolina (approval #18-203).

For behavior studies, male *Kif3a*^+/-^ (B6.129-Kif3a*^tm1Gsn^*/J, Stock No: 003537; Jackson Laboratories, Bar Harbor, ME) and female C57BL/6J mice were obtained, housed, and mated as described above to produce *Kif3a*^+/+^ and *Kif3a*^+/-^ offspring. Generation of *Kif3a^-^*^/-^ mice was avoided as this mutation is embryonically lethal. All dams were singly housed in clean cages on E15. Alcohol- and vehicle-treated litters (see below) were housed with their dams, culled to a maximum of 8 pups/litter at postnatal day (PD) 3, weighed, ear marked, and tail-clipped for genotyping on PD14, and left undisturbed until weaning at PD28 when they were housed in same sex groups with their littermates. All the mice from each litter were tested on behavioral experiments conducted between PD28–33 by an experimenter who was unaware of the prenatal treatments and *Kif3a* genotype.

### Neurulation-stage alcohol exposure (NAE)

On E9.0, pregnant dams were administered two doses of ethanol (25% vol/vol ethyl alcohol, Pharmaco-Aaper, Brookfield, CT, at a dose of 2.9 g/kg) in Lactated Ringer’s solution 4 hr apart via intraperitoneal (i.p.) injection. These mice were designated as the NAE group. A separate group of mice were administered an equal volume of vehicle (1.5 ml/100 g body weight). This model of alcohol exposure has been previously shown to result in maternal blood alcohol concentrations of ∼400 mg/dl (27). For molecular studies, dams were humanely sacrificed via CO_2_ followed by cervical dislocation either 6, 12, or 24 hr after the first ethanol injection. Embryos were dissected out and placed into cold RNase-free Dulbecco’s saline solution. For all assays, embryos were stage-matched based on somite number (E9.25: 21-22 somites, E9.5: 24-25 somites, E10: 30-31 somites). A maximum of two embryos per litter were used in each assay to minimize litter effects.

### Embryo immunohistochemistry

The chorion and amnion were removed from all embryos and placed into 4% paraformaldehyde for ∼72 h. Stage-matched embryos were then processed for paraffin-embedding on a Leica tissue processing station and sectioned at 7 µm on a microtome. A total of 6-10 sections per embryo, encompassing the rostroventral neural tube (RVNT), were processed for immunohistochemistry. Briefly, paraffin residue was dissolved in xylenes and sections were rehydrated in ethanol washes. Slides were quenched in 10%H_2_O_2_/90% methanol for 10 min and incubated for 30 min in 10% Citra Plus Buffer (BioGeneX, Fremont, CA) in a steam chamber. The slides were then blocked in normal goat serum/Triton-X/bovine serum albumin blocking solution for 1 hr and incubated overnight with primary antibody (Arl13b, 1:500, NeuroMabs, University of California-Davis) at 4°C for 24 h. Slides were then incubated with secondary antibody (anti-mouse Alexafluor 488, 1:1000, Thermofisher, Waltham, MA) for 2 hr at room temperature and cover slipped with Vectashield HardSet Antifade Mounting Medium with DAPI (Vector, Burlingame, CA).

### Confocal imaging and image quantification

Cilia were imaged on a Zeiss 880 confocal microscope with a 40x oil lens. Image stacks were obtained at a step of 0.46 µm between images. Stacks were then compressed into a single image containing a depth color code with Fiji (ImageJ) software (28). Cilia within the known volume of the RVNT were counted with the Cell Counter plug-in and expressed as number of cilia per 100 µm^3^. RVNT volume and cell layer width were determined using images of the same sections taken with a 10x lens. The DAPI-labeled cell layer was traced to obtain volumetric measurements and the cell layer width was measured at multiple points within each RVNT (2 measurements per slide) using Fiji software.

### Gene expression assays

The RVNT of stage-matched embryos were dissected and placed into lysis buffer and stored at -80°C until processing. RNA was isolated from the supernatant using the RNeasy Plus Micro Kit (Qiagen, Valencia, CA). Nucleic acid concentration and quality were determined using the Qubit 3.0 fluorometer and Nanodrop 2000 (Thermofisher, Waltham, MA). Generation of cDNA used a consistent starting amount of RNA across all embryos (100 ng). Multiplex qRT-PCR was used to determine alcohol-induced gene expression changes (n = 6-10 embryos/treatment). Taqman probes (Invitrogen) and Taqman Multiplex PCR mix (Applied Biosystems) were used to determine expression changes in the following genes: *Shh* (Mm00436528), *Gli1* (Mm00494654), *Gli2* (Mm01293117), *Ccdn1* (Mm00432359), *Ccdn2* (Mm00438071), *Fgf15* (Mm00433278), *Cep41* (Mm00473478), *Hap1* (Mm00468825), *Rilpl2* (Mm01199587)*, Evc* (Mm00469587)*, Nek4* (Mm00478688), and *Dpcd* (Mm00620237). For all assays, *18s* (Mm03928990) or *Pgk1* (Mm000435617) were used as reference genes. Reference genes were confirmed to be unaffected by prenatal treatment in separate embryos prior to use in these experiments. However, *18s* expression showed high baseline variability in the E9.25 embryos, leading to the use of *Pgk1* as a reference gene for this time point. All reactions were run in triplicate and amplicon specificity was confirmed with gel electrophoresis.

### Western Blot

RVNT’s of each litter (n = 5-10 samples per tube) were pooled and lysed in RIPA buffer with 1X Halt protease and phosphatase inhibitors (Thermofisher, Waltham, MA) in order to obtain sufficient protein for western blot analyses. For E9.25, 5 vehicle and 6 NAE litters were used, and for E9.5, 6 vehicle and 7 NAE litters were used. Protein concentrations were determined with the Micro BCA kit (Thermofisher, Waltham, MA) and colorimetrics measured at 540 nm on a spectrophotometer. 15 µg of sample with 4X Laemmli sample buffer were added to each lane of a tris-glycine gel in 1X TGX running buffer and run at 150V for 45 min. Protein weights were determined with the Dual Color protein ladder (Bio-Rad, Hercules, CA). Proteins were transferred to a membrane using the iBlot2 system and incubated overnight at 4°C with primary antibody (anti-Gli3, 1:500, AF3690, R&D Systems, Minneapolis, MN). Membranes were then incubated in secondary antibody (anti-goat Alexafluor 680, 1:10,000, Thermofisher, Waltham, MA) for 1 hr at room temperature and imaged on a Licor Odyssey scanner. GAPDH (1:3000, 14C10, Cell Signaling, Danvers, MA) was used as an internal loading control. The relative intensities of the Gli3 protein bands (Gli3^FL^ [∼190 kDa] and Gli3^Rep^ [∼83 kDa]) were analyzed with ImageJ Gel Analyzer software and normalized to internal control bands. The ratio of Gli3^FL^: Gli3^Rep^ forms was calculated and expressed as a percentage of total Gli3.

### Adolescent behavioral procedures

All the mice from each litter generated from the mating of a *Kif3a*^+/-^ male and C57BL/6J female (n = 16 litters vehicle-exposed and 19 litters NAE) were tested on behavioral experiments conducted between PD28–33 by an experimenter unaware of the prenatal treatments and *Kif3a* genotype. The experiments were conducted in the UNC Behavioral Phenotyping Core during the light portion of the 12:12hr light:dark schedule. All mice were tested in the following testing order: rotarod trials 1-3; elevated plus maze (EPM); open-field; rotarod trials 4 and 5. The EPM was tested before the open field to minimize potential carry-over effects, as the EPM is more sensitive to the effects of prior testing history (29, 30). Technical issues during testing resulted in two male animals being excluded from the EPM and one male from open field analyses.

The rotarod (Ugo-Basile, Stoelting Co., Wood Dale, IL) measured the latency to fall off or rotate around the top of a dowel which progressively accelerated from 3 rpm to 30 rpm during the maximum of a 5-min test, conducted during three repeated trials on the first day of testing and two repeated trials on the second day of testing. Each trial was separated by ∼45 sec. The EPM was 50 cm above the floor and contained two open arms (30 cm length, 220 lux) and two closed arms (20 cm high walls, 120 lux). During the 5-min test, an observer recorded the number of entries and time spent in each of the arms. These data were used to calculate the percent of open arm time [(open arm time/total arm time) × 100], the total arm entries, and the percent of open arm entries [(open arm entries/total arm entries) × 100]. The open field (41 × 41 × 30 cm) was illuminated (120 lux at the edges, 150 lux in the center), housed within a sound-attenuated chamber and equipped with upper and lower grids of photobeams for the detection of horizontal and vertical activity and position of the mouse (Versamax System: Accuscan Instruments, Columbus, OH). The open field was conducted over 60 min and the primary measures were horizontal activity, total distance traveled, time spent in the center of the chamber, and distance traveled in the center. Number of rears and rotations were also recorded. Open field data were analyzed as four separate 15-min epochs (i.e. min 0-15, min 16-30, min 31-45, and min 46-60), based on prior findings that the initial minutes of the open field test are most sensitive to developmental drug exposure (11, 31).

### Adolescent brain measurements

Following completion of behavioral tasks on PD37, mice were deeply anesthetized with tribromoethanol anesthesia and transcardially perfused with 1X PBS followed by 10% formalin. Brains were removed and stored in formalin for at least one week. Prior to paraffin embedding, fixed whole brains were cleared by rocking at room temperature in 1X PBS for one week and 70% ethanol for one week (each solution changed daily). Brains were then cut into two sections, the frontal lobes and cerebellum, and paraffin-embedded on a Leica tissue processing station. Tissue was then embedded in paraffin block and sectioned at 10 µm on a microtome. Every 5^th^ slide (containing 3-4 sections per slide) throughout the entirety of the cortex was stained using cresyl violet. One section per slide was imaged and the ventricles traced using Fiji/ImageJ (28). Total area of the ventricles were summed per section and averaged across the entire brain. For midline brain width and medial height, lines were drawn across images of one section of each slide throughout the cortex and measured in Fiji. Midline brain width was measured at the widest part of the cortex (dorsal/ventral bregma ∼3.0) and height was measured as close to the midline as possible (medial/lateral bregma 0.25); measurements were averaged across the entire brain.

### Statistical analyses

Cilia number, RVNT volume, and Western blot band intensity were analyzed using unpaired *t*-tests comparing prenatal treatment (NAE vs. vehicle) at each time point with Welch’s correction when necessary. The Benjamini-Hochberg correction was used if multiple unpaired *t*-tests were run within an experiment (false discovery rate = 0.10) (32). Raw *p*-values shown in tables and in the text remain statistically significant following correction unless otherwise noted. Multiplex qRT-PCR gene expression data are expressed as log2 fold change calculated using the 2^(-ΔΔCT)^ method (33). Data were analyzed using either unpaired *t*-tests with corrected *p*-values (cilia-related genes) or two-way ANOVAs (Treatment × Time Point) with corrected *t*-tests run as *post hocs* for analyses with significant interactions (Shh path genes, Gli3, cell cycle genes). For behavior assays, litter means were analyzed for each treatment and genotype to control for between litter effects (34). Males and females were analyzed separately, because our previous studies demonstrated sex differences on these measures (12, 35). EPM data were analyzed using two-way ANOVAs (treatment × genotype) and open field and rotarod data were analyzed with three-way ANOVAs (treatment × genotype × time bin or trial). For the behavioral analysis, we predicted, *a priori*, that any effects of NAE or *Kif3a*^+/-^ would be exaggerated when these treatments were combined, in other words, that the NAE *Kif3a*^+/-^ mice would have the largest difference from the vehicle-treated *Kif3a*^+/+^ mice. Ventricle area measurements were analyzed using a three-way ANOVA (treatment × genotype × bregma), while width and height measurements were analyzed with two-way ANOVAs (treatment × genotype). *Post hoc* analyses were conducted when appropriate. Statistical differences were considered significant at an adjusted *p*-value threshold of 0.05.

## Results

### NAE downregulates expression of the Shh pathway in the RVNT

Previous work has demonstrated that the shape and size of the ventricles, pituitary, and hypothalamus are altered following NAE (11–14), suggesting that during neurulation the RVNT (from which these structures are derived) is particularly vulnerable to alcohol. Dysregulation of Shh signaling within the RVNT is likely a mechanism for structural changes to midline brain regions following NAE, however, this hypothesis has never been tested (36). To determine whether NAE alters the Shh pathway in the RVNT, we administered two doses of 2.9 g/kg alcohol during mid-neurulation (E9.0) to model binge-like alcohol exposure. We collected the RVNT of stage-matched embryos 6, 12, and 24 hr later (E9.25, E9.5, and E10, see methods for somite numbers) and analyzed *Shh, Gli1,* and *Gli2* gene expression. For each gene, two-way ANOVAs were run using the factors of Treatment × Time point (statistics in Table 1), with *post hoc t*-tests between NAE and Veh groups performed for each time point in the event of a significant interaction. First, we found main effects of Treatment and Time Point and a significant interaction for *Shh* gene expression. *Post hocs* revealed that NAE significantly downregulated *Shh* at both the 6 and 12 hr time points (*t*(14) = 2.741, *p* = 0.0159 and *t*(14) = 3.608, *p* = 0.003, respectively; Fig 1A; Table 1), extending previously reported reductions in Shh following gastrulation-stage alcohol (9, 10) and demonstrating that NAE also impairs Shh levels in the RVNT, but is independent of concurrent apoptosis in this region (5). Following Shh activation, Gli1 is rapidly upregulated, whereas Gli2 is constitutively expressed. However, without Shh present, Gli2 is cleaved and tagged within the primary cilium for degradation. Activation of Smoothened (Smo) by Shh deactivates this cilia-mediated cleavage of Gli2 and allows it to be shuttled outside of the cilia to activate downstream genes (37). A main effect of treatment was also found for Gli1, with NAE embryos showing reduced *Gli1* expression compared to controls (Table 1, Fig 1A). No effects of NAE were found for *Gli2* expression. In addition, we measured the relative amounts of the two forms of Gli3 protein 6, 12, and 24 hr following the beginning of NAE. In the absence of Shh and with normal cilia functioning, Gli3 is found predominantly in the cleaved repressor form (Gli3^Rep^). Upon Shh pathway activation, the balance of Gli3^Rep^ to the full-length activator form (Gli3^FL^) is tipped slightly to the activator form to allow Gli1 and Gli2 to proceed with normal gene transcription, including cell cycle genes such as the cyclin family, as well as having a positive feedback effect on Shh itself (38). A two-way ANOVA revealed a significant Treatment × Time interaction (statistics in Table 1). *Post hoc t*-tests found that NAE led to a significantly shifted ratio at the 12 hr time point, with a higher percentage of Gli3^Rep^ compared to vehicle treatment (65.27% Gli3^Rep^ in NAE vs. 54.7% Gli3^Rep^ in vehicle-treated; Fig 1B; *t*_(11)_ = 2.67, *p* = 0.022).

**Fig 1.**
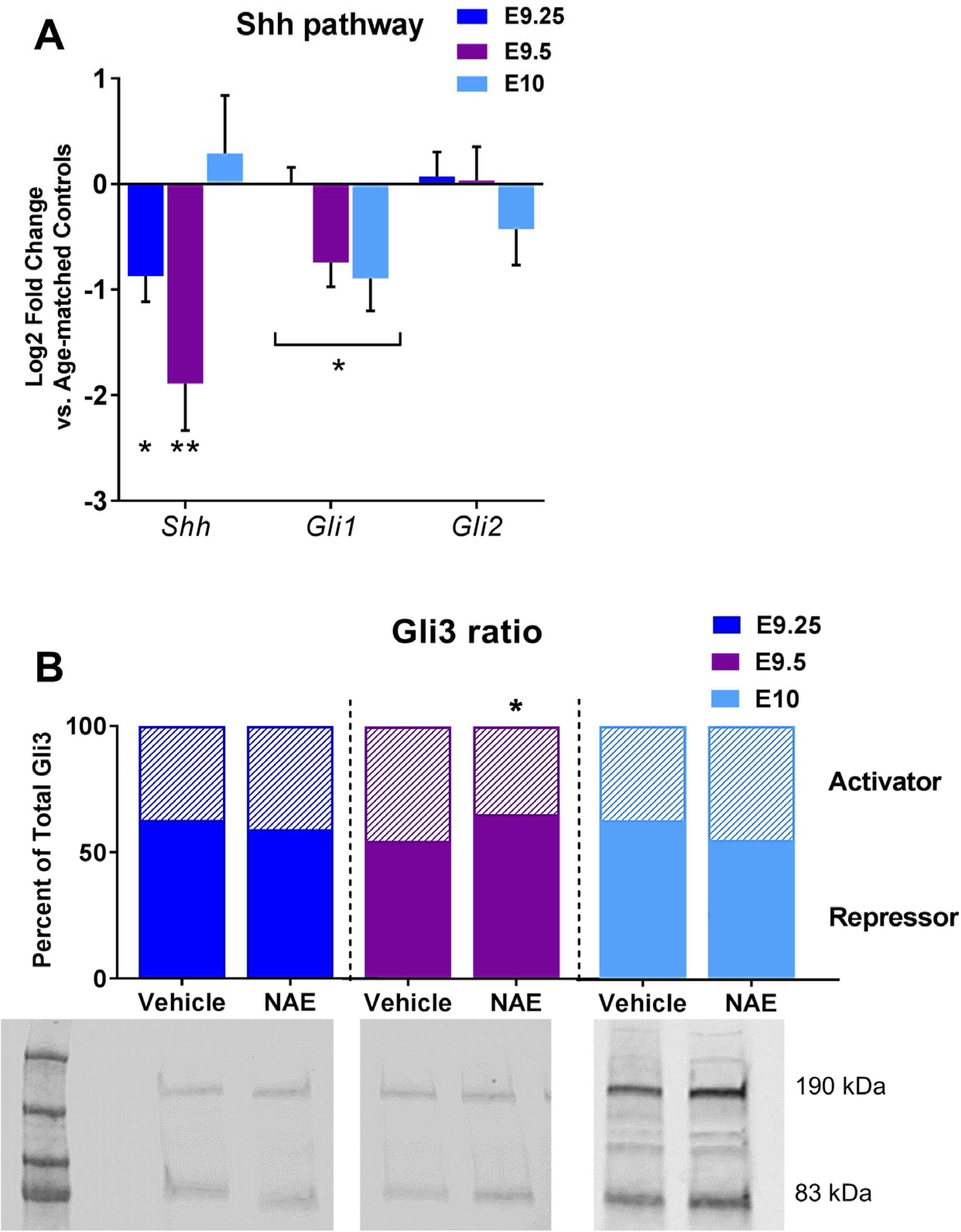
Neurulation-stage alcohol exposure (NAE) decreases *Shh* pathway activation in the RVNT. A) NAE resulted in downregulation of *Shh* and *Gli1* within 24 hr post-exposure (n = 7-10 embryos per group). *Shh* expression was significantly reduced 6 and 12 hr after NAE. Expression of *Gli2* did not differ. Brackets indicate a significant main effect of prenatal treatment. Gene expression data are the log2 fold change compared to somite-matched embryos from the vehicle group (expressed as 0 on the graph) ± SEM. B) The relative percentage of Gli3^Rep^ to Gli3^FL^ was significantly higher in NAE embryos (n = 7 litters) compared to the vehicle group (n = 6 litters) 12 hr after exposure. Data are expressed as the average Gli3^FL^ or Gli3^Rep^ band density expressed as a percentage of total Gli3. Below are representative bands of Gli3^FL^ (190 kDa) and Gli^Rep^ (83 kDa). Full blot and loading control GAPDH (37 kDa) can be seen in S1 Fig. * = *p* < 0.05, ** = *p* < 0.01.

**Table 1.** Statistical results for Shh pathway and cell cycle genes. Gli3 results refer to protein quantification of Gli3^Rep^:Gli3^FL^ ratio. Significant (*p* < 0.05) results are listed in bold.

| Gene | Associated protein | Function | Log2 Fold Change | Two-way ANOVA<br>(Treatment x Time Point) |
| --- | --- | --- | --- | --- |
| *Shh* | Sonic Hedgehog | Morphogen,<br>neural<br>patterning | E9.25: -0.87<br>E9.5: -1.89<br>E10: 0.29 | **<u>Treatment:</u> $F(1,43) = 6.637, p = 0.014$**<br>**<u>Time Point:</u> $F(2,43) = 3.845, p = 0.029$**<br>**<u>Interaction:</u> $F(2,43) = 3.916, p = 0.027$** |
| *Gli1* | GLI Family Zinc Finger 1 | Transcription<br>factor | E9.25: 0.01<br>E9.5: -0.74<br>E10: -0.89 | **<u>Treatment:</u> $F(1,43) = 6.814, p = 0.012$**<br><u>Time Point:</u> $F(2,43) = 1.956, p = 0.154$<br><u>Interaction:</u> $F(2,43) = 2.171, p = 0.126$ |
| *Gli2* | GLI Family Zinc Finger 2 | Transcription<br>factor | E9.25: 0.698<br>E9.5: 0.035<br>E10: -0.43 | <u>Treatment:</u> $F(1,44) = 0.121, p = 0.73$<br><u>Time Point:</u> $F(2,44) = 0.452, p = 0.639$<br><u>Interaction:</u> $F(2,44) = 0.662, p = 0.521$ |
| *Ccnd1* | Cyclin D1 | Cell cycle | E9.25: -0.387<br>E9.5: -0.62<br>E10: -0.48 | **<u>Treatment:</u> $F(1,43) = 12.735, p = 0.001$**<br><u>Time Point:</u> $F(2,43) = 0.232, p = 0.794$<br><u>Interaction:</u> $F(2,43) = 0.218, p = 0.805$ |
| *Ccnd2* | Cyclin D2 | Cell cycle | E9.25: -0.3288<br>E9.5: -0.29<br>E10: -0.94 | **<u>Treatment:</u> $F(1,43) = 9.772, p = 0.003$**<br><u>Time Point:</u> $F(2,43) = 1.7, p = 0.195$<br><u>Interaction:</u> $F(2,43) = 1.676, p = 0.199$ |
| *Fgf15* | Fibroblast Growth Factor 15 | Cell growth | E9.25: -1.107<br>E9.5: -0.25<br>E10: 0.038 | **<u>Treatment:</u> $F(1,43) = 5.981, p = 0.019$**<br>**<u>Time Point:</u> $F(2,43) = 3.917, p = 0.027$**<br><u>Interaction:</u> $F(2,43) = 2.272, p = 0.115$ |
| Gli3 | GLI Family Zinc Finger 3 | Transcription<br>factor | | <u>Treatment:</u> $F(1,26) = 0.0974, p = 0.756$<br><u>Time Point:</u> $F(2,26) = 0.300, p = 0.743$<br>**<u>Interaction:</u> $F(2,26) = 3.42, p = 0.048$** |

The shift in Gli3 to favor the repressor form at this time point coincides with significant reductions in *Shh* and *Gli1* gene expression, as well as smaller RVNT volumes in NAE embryos (Fig 1). The ratio of Gli3^FL^:Gli3^Rep^ in the RVNT did not significantly differ from control levels 6 or 24 hr post-NAE (Fig 1B). No difference in total amount of Gli3 was observed between treatment groups at any time point and the amount of Gli3^Rep^ did not significantly differ between the E9.25, E9.5, or E10 vehicle-treated groups (F_(2,14)_ = 0.967, *p* = 0.401). Changes to Shh pathway transduction could alter the developmental trajectory of craniofacial and CNS regions derived from the RVNT by affecting normal cell proliferation, resulting in the midline defects observed in mice following NAE.

### NAE downregulates cell cycle gene expression and decreases RVNT volume

One of the outputs of the Shh signaling pathway is the expression of cell cycle genes. Based on the observed decreases in *Shh* and *Gli1* gene expression and altered Gli3^FL^:Gli3^Rep^ following NAE, we hypothesized that NAE would also lead to altered expression of genes involved in regulation of cell proliferation. To assess changes in Shh-mediated gene expression, we analyzed three cell proliferation genes downstream of Shh, Cyclin D1 (*Ccnd1*), Cyclin D2 (*Ccnd2*), and Fibroblast growth factor 15 (encoded by *Fgf15*; homologous to *Fgf19* in humans), in the RVNT 6, 12, and 24 hr following NAE. Ccnd1 and Ccnd2 regulate progression of the G1 phase of the cell cycle (39) and are regulated by the Shh pathway Gli family of transcription factors (40, 41). Furthermore, Ccnd1 is important for expansion of ventral neural precursors in the early mouse diencephalon (42). In accordance with the alcohol-induced reductions in *Shh*, significant main effects of treatment were found for *Ccnd1* and *Ccnd2* (Fig 2A, Table 1), with NAE embryos displaying reduced expression compared to vehicle-treated embryos. Fgf15 is dependent on Shh signaling and largely overlaps with Gli1 in expression patterns early in development (42, 43). Functionally, Fgf15 promotes cell cycle exit and differentiation of neuronal precursors (44). Main effects of both treatment and time point were seen for Fgf15 expression (Fig 2A; Table 1), however the NAE-induced reduction in *Fgf15* was most pronounced on E9.25.

**Fig 2.**
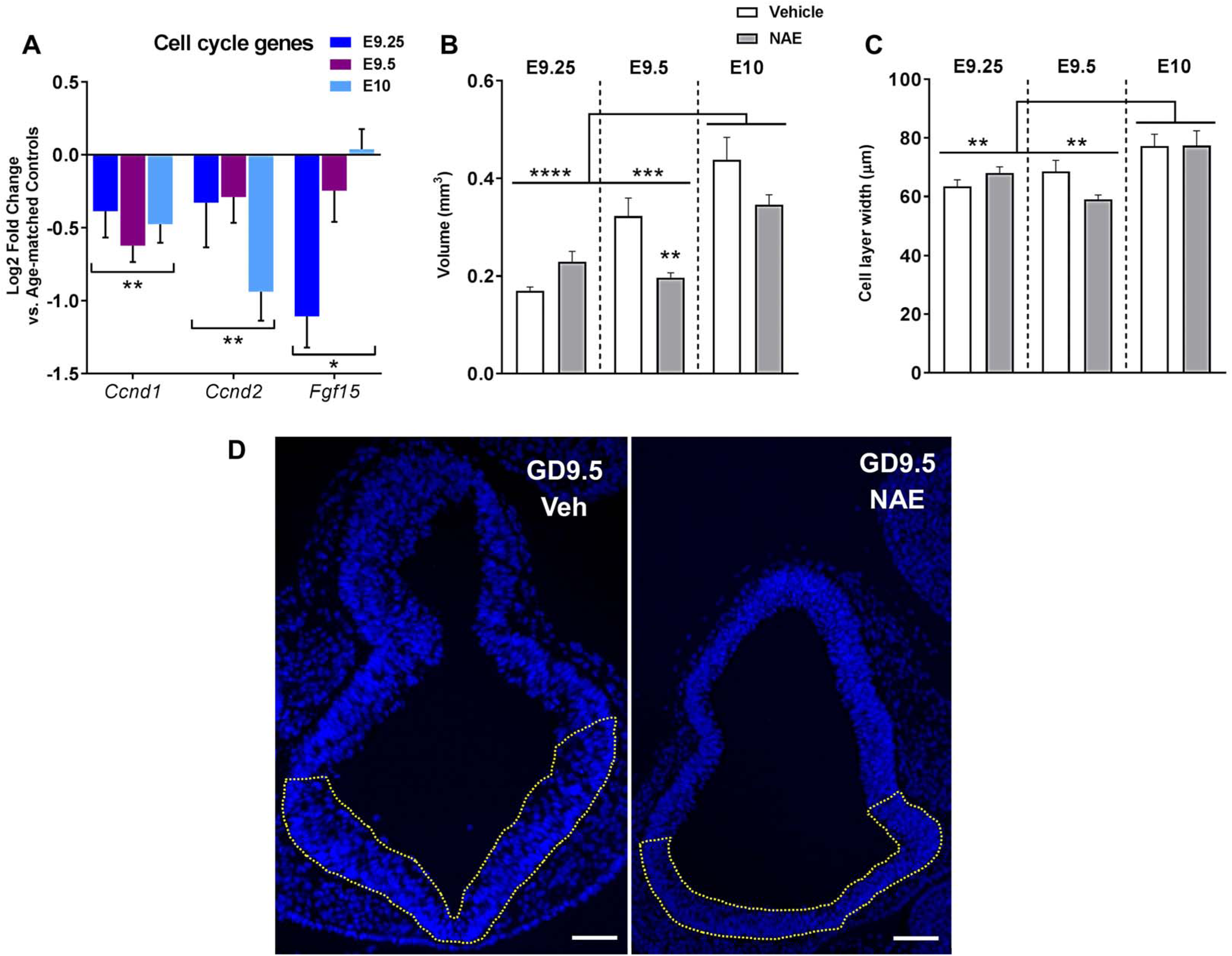
NAE decreases expression of Shh-mediated cell proliferation genes in the RVNT and reduces RVNT volume. A) NAE reduced expression of key Shh-mediated genes important for cell proliferation processes (n = 6-10 embryos per group) within 24 hr post-exposure. Brackets indicate main effects of prenatal treatment (Table 1). Gene expression data are the log2 fold change compared to the vehicle group (expressed as 0 on the graph) ± SEM. B) RVNT volume is significantly reduced in NAE embryos (filled bars, n = 7) compared to vehicle-treated embryos (open bars, n= 9) 12 hr after exposure, but did not differ at the other two time points. E10 RVNTs were also significantly larger than both E9.25 and E9.5. C) Cell layer width did not differ based on prenatal treatment but did increase over time, with E10 embryos exhibiting wider RVNT cell layers compared to both E9.25 and E9.5 embryos. D) Representative images of the RVNT of E9.5 vehicle-treated vs. NAE embryos taken with a 10x lens. Scale bar = 100 µm. * = p < 0.05, ** = *p* < 0.01, *** = p < 0.001, **** = p < 0.0001, all values expressed as mean + SEM.

Given the disruption of cell cycle-related gene expression, we next analyzed the volume of the RVNT 6, 12, and 24 hr after NAE to examine how the size of this region changed over time and in response to alcohol (Fig 2B). Images taken from the RVNT were obtained at 40x for volumetric analyses; section thickness was also measured at 40x. A significant time (hours post-exposure) × treatment interaction was observed (*F*_(2,39)_ = 6.164, *p* = 0.0047), as well as main effects of both time (*F*_(2,39)_ = 22.08, *p* < 0.0001) and treatment (*F*_(2,39)_ = 5.018, *p* = 0.031). As expected, the volume of the RVNT increased across time, with E10 embryos having significantly larger RVNTs compared to both E9.25 (*p* < 0.0001) and E9.5 (*p* = 0.0002), demonstrating the exceptional growth of the neural tube over this period. There was a trend towards the RVNT of E9.5 embryos being larger than E9.25 (*p* = 0.065), however this relationship was not statistically significant. Importantly, RVNT volume was significantly smaller in NAE compared to vehicle-treated embryos (0.197±0.01 mm^3^ vs. 0.3224±0.04 mm^3^) on E9.5 (*p* = 0.0085) and tended to be smaller on E10 as well, though this difference did not reach statistical significance (*p* = 0.1). The thickness of the RVNT cell layer was also assessed, however while the cell layer did grow thicker across time (*F*_(2,39)_ = 9.142, *p* = 0.0006), there was no significant effect of treatment (Fig 1C). E10 embryos had increased widths compared to both E9.25 (*p* = 0.0049) and E9.5 (*p* = 0.0012). The increase in cell layer width corresponds to the larger overall volume at E10. The rapid changes in RVNT volume across the first 24 hr post-alcohol exposure indicate that significant structural changes are taking place and encourages further investigation into all factors by which alcohol could affect growth, including alterations to cell proliferation. Additionally, while NAE-induced dysmorphologies are evident later in development (11–14), no overt gross anatomical differences were observed between vehicle and NAE-treated embryos at E9.5 (S2 Fig). Thus, alcohol-related changes in tissue volume and shape are likely subtle at this time point and region- and timing-specific. Even though the volumetric changes induced by NAE were transient, a brief period of growth inhibition within this region of the neural tube during such an important and dynamic period of early development could have long-lasting consequences on face and brain structure.

### NAE alters expression of key cilia-related genes, but not cilia density in the RVNT

Shh transduction occurs within the axoneme of primary cilia, nearly ubiquitous structures of mammalian cells that play a particularly important role during embryonic development. Dysregulation of RVNT primary cilia poses a risk to the normal development of brain regions that arise from this area of the neural tube, namely midline structures such as the hypothalamus, septum, pituitary, and preoptic area (45), many of which have been shown to be altered by NAE Additionally, midline anomalies such as hypertelorism are a hallmark of genetic ciliopathies and have also been reported in a subset of patients with heavy prenatal alcohol exposure (4), suggesting an overlapping etiology between ciliopathies and prenatal alcohol.

Based on the reduction in *Shh* and downstream cell cycle gene expression, we hypothesized that we might find a coinciding NAE-induced change in either 1) the number of primary cilia or 2) cilia-related genes in the RVNT. First, we examined the direct impact of NAE on primary cilia in the RVNT by analyzing cilia density. Primary cilia were labeled for the cilia-specific small GTPase Arl13b at 6, 12, or 24 hr following NAE (Fig 3A) and the number of cilia within the known volume of the RVNT were analyzed using 3D image stacks from confocal images. A significant time (hours post-exposure) × treatment interaction was found (*F*_(2,39)_ = 3.642, *p* = 0.036), as well as a main effect of time (*F*_(2,39)_ = 18.93, *p* < 0.0001) (Fig 3B). No main effect of treatment was observed. No significant differences were seen between vehicle and NAE embryos at any time point, though there was a trend towards an increase in cilia density at the E9.5 (12 hr post-exposure) time point (*p* = 0.087). The density of cilia in the RVNT significantly decreased over time, as there was a higher density of cilia at E9.25 compared to both E9.5 and E10 (*p* < 0.0001 for both time points). This reduction in density is likely due to the substantial growth of the neural tube during neurulation (Fig 2B-C), however the lack of a treatment effect precludes alcohol-induced effects on cilia number as the mechanism of action of reduced cell cycle gene expression.

**Fig 3.**
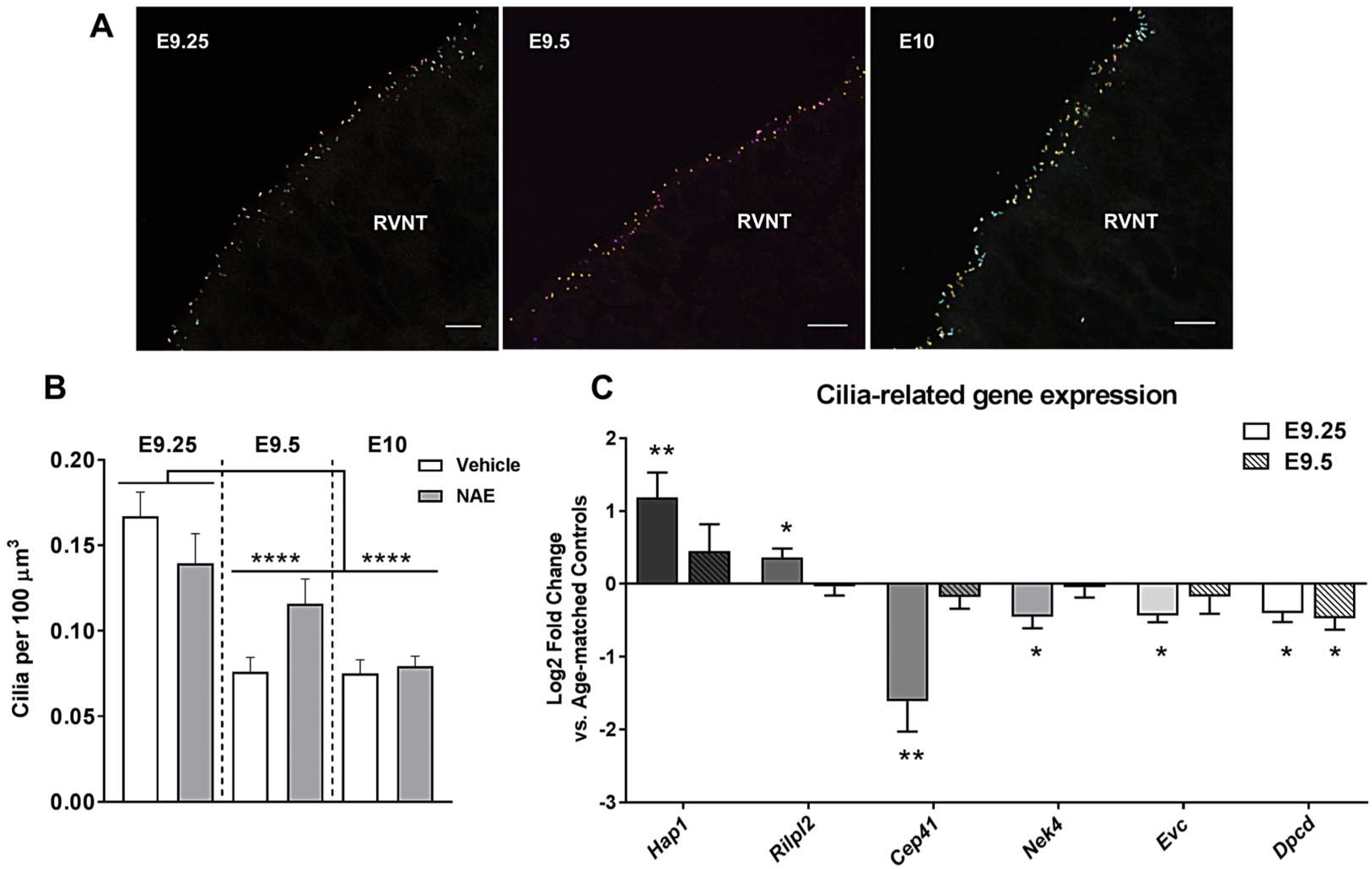
NAE alters expression of cilia-related genes in the RVNT but does not alter cilia density. A) Primary cilia in the RVNT were labeled with an anti-Arl13b antibody and stacks were compressed to visualize cilia throughout the depth of the tissue in a single image. Cilia in the images are pseudo-colored, scale bar = 10 µm. B) Cilia density did not differ at any of the time points, but did decrease across time points, as E9.5 and E10 had lower cilia density compared to E9.25 (n = 6-9). ** = *p* < 0.01, **** = p < 0.0001, all values are expressed as mean + SEM. All gene expression data are the log2 fold change compared to the vehicle group (expressed as 0 on the graph) ± SEM. C) NAE resulted in significant changes to expression levels of genes related to ciliogenesis and post-transcriptional modifications to ciliary tubulin, as well as genes implicated in genetic ciliopathies 6 hr after exposure. At 12 hr post-exposure, all genes had returned to baseline expression levels except for *Dpcd*, which remained significantly downregulated. E10 expression was not evaluated based on the return to baseline observed for almost all genes at E9.5. Encoded proteins associated with these genes can be found in Table 2. For all genes, n = 6-8 embryos per group. * = *p* < 0.05, ** = *p* < 0.01. Data are expressed as log2 fold change compared to the vehicle group (expressed as 0 on the graph) ± SEM.

**Table 2.**
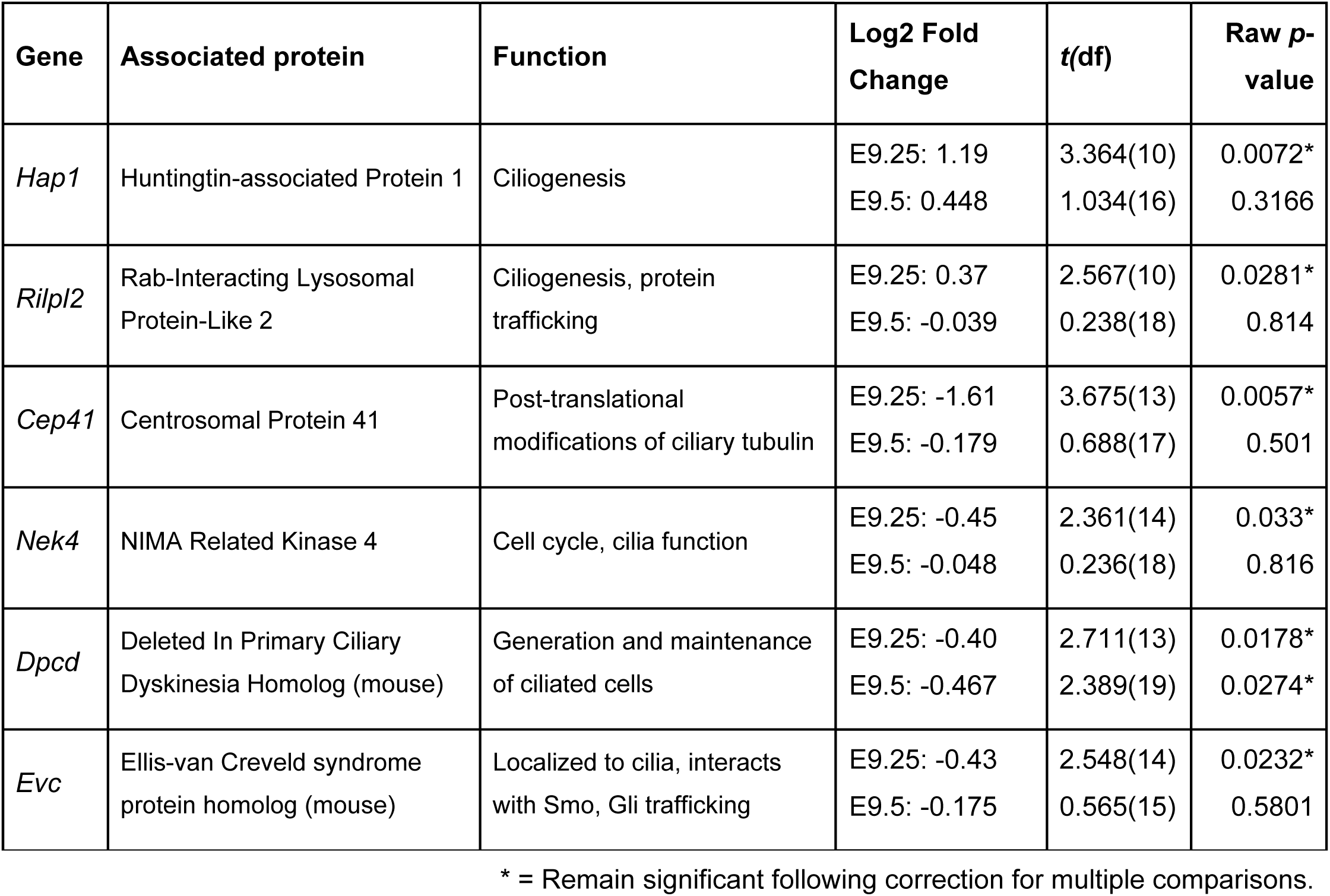
Statistical results for cilia-related genes.

| Gene | Associated protein | Function | Log2 Fold Change | t(df) | Raw p-value |
| --- | --- | --- | --- | --- | --- |
| *Hap1* | Huntingtin-associated Protein 1 | Ciliogenesis | E9.25: 1.19<br>E9.5: 0.448 | 3.364(10)<br>1.034(16) | 0.0072*<br>0.3166 |
| *Rilpl2* | Rab-Interacting Lysosomal Protein-Like 2 | Ciliogenesis, protein trafficking | E9.25: 0.37<br>E9.5: -0.039 | 2.567(10)<br>0.238(18) | 0.0281*<br>0.814 |
| *Cep41* | Centrosomal Protein 41 | Post-translational modifications of ciliary tubulin | E9.25: -1.61<br>E9.5: -0.179 | 3.675(13)<br>0.688(17) | 0.0057*<br>0.501 |
| *Nek4* | NIMA Related Kinase 4 | Cell cycle, cilia function | E9.25: -0.45<br>E9.5: -0.048 | 2.361(14)<br>0.236(18) | 0.033*<br>0.816 |
| *Dpcd* | Deleted In Primary Ciliary Dyskinesia Homolog (mouse) | Generation and maintenance of ciliated cells | E9.25: -0.40<br>E9.5: -0.467 | 2.711(13)<br>2.389(19) | 0.0178*<br>0.0274* |
| *Evc* | Ellis-van Creveld syndrome protein homolog (mouse) | Localized to cilia, interacts with Smo, Gli trafficking | E9.25: -0.43<br>E9.5: -0.175 | 2.548(14)<br>0.565(15) | 0.0232*<br>0.5801 |

\* = Remain significant following correction for multiple comparisons.

We further investigated primary cilia homeostasis and stability in the RVNT through analysis of six cilia-related genes 6 or 12 hr following NAE: *Hap1*, *Rilpl2*, *Cep41*, *Nek4*, *Evc*, and *Dpcd.* These time points were focused on since almost all genes returned to baseline expression after 12 hr (Table 2, Fig 3C). These genes were chosen due to their known roles in important ciliary processes including ciliogenesis, ciliary protein trafficking, or cilia stability through maintenance of post-translational modification and/or genes that have mutations associated with human genetic ciliopathies. Based on our *a priori* hypothesis that cilia-related genes would be most affected at the 6 hr time point (E9.25), each time point was analyzed separately. At E9.25, NAE upregulated two of the genes, *Hap1* and *Rilpl2* (1.19 and 0.37 log2 fold change, respectively), in the RVNT (Table 2; Fig 3C). However, expression levels of these genes returned to baseline by E9.5 (Table 2, Fig 3C). The other four genes (*Cep41*, *Nek4*, *Evc*, and *Dpcd*) showed marked downregulation 6 hr following NAE (E9.25) (Table 2, Fig 3C). *Dpcd* remained significantly downregulated (-0.467 log2 fold change; Table 2, Fig 3C) 12 hr after NAE; however, expression of *Cep41*, *Nek4*, and *Evc* had returned to control levels. These genes have prominent roles related to cilia function, stability, and ciliogenesis. The NAE-induced dysregulation of these genes precedes and coincides with the downregulation of important morphogenic and cell proliferation pathways as well as reductions in RVNT volume.

### Partial knockdown of cilia gene *Kif3a* interacts with NAE to affect adolescent behavior

To further test the hypothesis that disruption of normal cilia function is a mechanism for the consequences of NAE, we utilized a transgenic mouse strain with a partial deletion of *Kif3a*, a gene that codes for an intraflagellar transport protein. Full deletion of *Kif3a*, is a well-characterized ciliopathy model (24, 26, 46, 47), and if NAE interacts with cilia function, then *Kif3a* heterozygosity would both phenocopy and exacerbate NAE. We focused on a NAE effect that we have previously shown to be robust and reproducible, behavioral change during adolescence (11). Adolescent male and female mice exposed to E9.0 alcohol were tested on a battery of behavioral tasks, but since the largest behavioral changes occurred in males, we discuss behavior from male, rather than female, mice in detail (for female data refer to Fig S3B, S4A-D; S1 Table). Sex-specific effects of early gestational alcohol have been previously reported to emerge in adolescence and adulthood and likely result from postnatal sex differences, as opposed to embryonic effects of alcohol

We first examined motor coordination, using an accelerating rotarod performance across five repeated trials. Although rotarod performance for all mice improved from the first to the fifth trial (*F*_(4,192)_ = 28.4, *p* < 0.0001), NAE persistently impaired motor coordination, as revealed by a main effect of NAE, regardless of *Kif3a* genotype (*F*_(1,48)_ = 5.1; *p* = 0.028; Fig S3). Previous research has suggested that the etiology of cerebellar deficits caused by mid-stage NAE is likely alcohol-induced apoptosis in the rostral rhombic lip, from which the cerebellum arises (5). The lack of an NAE × genotype interaction indicates that primary cilia are not critical for these motor incoordination effects. While it is possible that alcohol earlier or later in development could have more direct effects on primary cilia-mediated mechanisms in the cerebellum, the middle of neurulation appears to be a critically sensitive period for cerebellar defects, as our previous research has suggested that alcohol administered on E7 (O’Leary-Moore & Sulik, *unpublished observations*) or E8 (11) has a less pronounced impact on motor coordination.

However, when we examined elevated plus maze (EPM) exploration, a two-way ANOVA found a significant genotype × treatment interaction (*F*_(1,49)_ = 6.9; *p* = 0.01) on total arm entries. Vehicle-treated *Kif3a*^+/-^ and NAE *Kif3a*^+/+^ mice were more active than were vehicle-treated *Kif3a*^+/+^ mice (*p* = 0.006 and *p* = 0.04, respectively) (Fig 4A), while NAE *Kif3a*^+/-^ had comparably high levels of activity but were not statistically different from vehicle-treated *Kif3a*^+/+^ mice (*p* = 0.061). This result replicates our previous findings on early NAE (11) and indicates that *Kif3a* heterozygosity phenocopies a locomotor hyperactivity caused by NAE. Supporting the hypothesis that NAE and *Kif3a*^+/-^ also affect anxiety-like behavior, there was a significant genotype × treatment interaction on the percentage of open arm time (*F*_(1,49)_ = 3.9; *p* = 0.05) (Fig 4B), but not the percent of open arm entries (Fig 4C). *Post ho*c analysis revealed that both vehicle-treated and NAE *Kif3a*^+/-^ mice spent a greater percentage of time on the open arms than did the vehicle-treated *Kif3a*^+/+^ mice (*p* < 0.001 and *p* = 0.01, respectively). Although NAE in *Kif3a*^+/+^ tended to phenocopy *Kif3a*^+/-^ on the open arm time, this effect was not significant in the *post hoc* analyses. To test whether NAE alone affects open arm time in the *Kif3a*^+/+^ mice and confirm our previous results using a two-sample *t*-test (11, 12), we ran this same statistical analysis (*t_(_*_20)_ = 3.0; *p* = 0.007; Fig 4B) and found that that early NAE increases the percent of time on the open arms. Overall, these results indicate that NAE and *Kif3a* heterozygosity phenocopy each other’s effects on the EPM.

**Fig 4.**
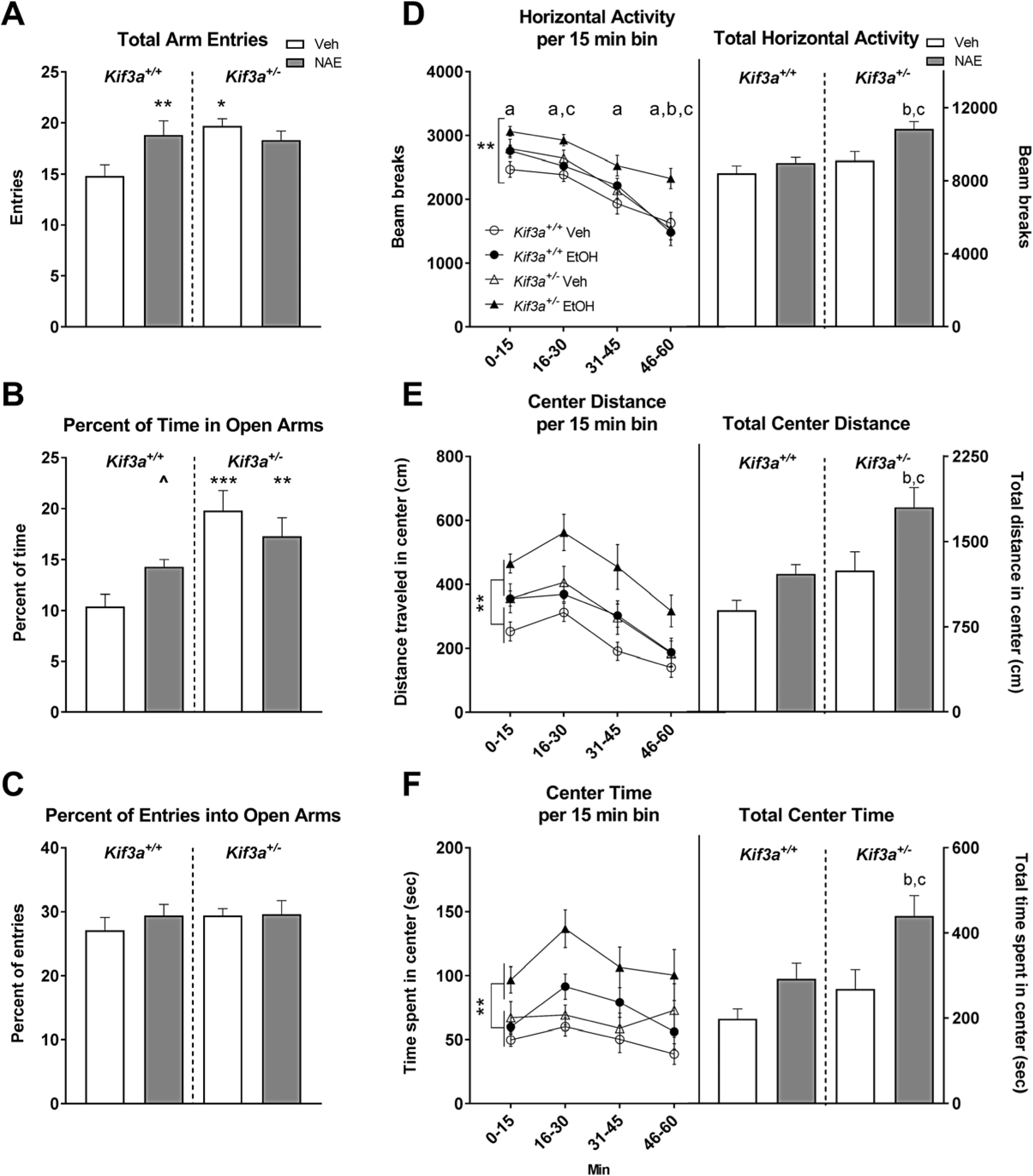
Partial loss of cilia motor transport gene *Kif3a* phenocopies and potentiates the effect of NAE on behavioral performance in adolescent male mice. A) Vehicle-treated *Kif3a*^+/-^ and NAE WT mice made more arm entries on the EPM than did vehicle-treated WT mice. B) Vehicle-treated and NAE *Kif3a*^+/-^ mice spent a greater percentage of time on the open arms vs. vehicle-treated WT mice. C) Percent of entries into the open arms was not significantly affected by genotype or treatment. D-F) Significant (*p* < 0.05) *post hoc*s are shown as letters on each graph. a: vs. vehicle-treated WT, b: vs. vehicle-treated *Kif3a*^+/-^, c: vs. NAE WT. D) NAE *Kif3a*^+/-^ mice were more active than vehicle-treated WT mice during all time bins and were more active than NAE WT mice during the 2^nd^ and 4^th^ bins and more active than vehicle-treated *Kif3a*^+/-^ mice during the final bin. Bracket = genotype × treatment interaction. NAE *Kif3a*^+/-^ mice had more total beam breaks compared to the NAE WT and vehicle *Kif3a*^+/-^ mice. E) NAE mice traveled further in the center of the open field compared to vehicle-treated mice. In addition, NAE *Kif3a*^+/-^ mice traveled further than NAE WT or vehicle *Kif3a*^+/-^ groups. Bracket = main effect of treatment. F) NAE animals spent more time in the center of the chamber compared to vehicle-treated mice across time, and NAE *Kif3a*^+/-^ mice spending more time in the center compared to both NAE WT and vehicle *Kif3a*^+/-^ mice. Bracket = main effect of treatment. For all groups, n’s = 11-15 litters, with same genotype littermates averaged into a single datum for each litter. For all graphs, * = *p* < 0.05, ** = *p* < 0.01, *** = *p* < 0.001. All data are shown as group means + SEM.

Since the EPM is a relatively brief exposure to a complex novel environment and our procedure did not assess habituation to the maze, we further measured exploratory behavior and activity in an open field test across a 1-hr period. A three-way mixed ANOVA revealed a significant genotype × treatment × time bin interaction on overall horizontal activity (*F*_(3,141)_ = 4.3; *p* = 0.006) (Fig 4D). *Post hoc* analysis revealed that NAE *Kif3a*^+/-^ mice were more active than vehicle-treated *Kif3a*^+/+^ mice during each of the 15-min time bins (*p* = 0.002, 0.001, 0.032, and 0.046, respectively), and were more active than NAE *Kif3a*^+/+^ mice during the 2^nd^ and the final 15-min time bins (*p* = 0.018 and 0.006, respectively). NAE *Kif3a*^+/-^ mice were also more active than vehicle-treated *Kif3a*^+/-^ mice during the final 15-min time bin (*p* = 0.013). Since the persistently high activity of the NAE *Kif3a*^+/-^ mice suggests impaired habituation, we calculated a habituation index, expressing data from each 15-min time bin as a percentage of the initial time bin (i.e. min 0-15). A significant 3-way interaction (genotype × treatment × time bin; *F*_(3,141)_ = 2.0; *p* = 0.039) revealed that while all groups reduced their activity over the hour, in the final time bin, NAE *Kif3a*^+/-^ mice were significantly less habituated than were the vehicle-treated *Kif3a*^+/-^ mice and the NAE *Kif3a*^+/+^ mice (*p* = 0.02 and 0.03, respectively). These groups had an initial bout of heightened activity, relative to the vehicle-treated *Kif3a*^+/+^ mice, but NAE *Kif3a*^+/-^ mice maintained a higher level of activity throughout the test.

There were significant main effects of NAE on several specific measures of activity, including repeated beam breaks (*F*_(1,47)_ = 4.1; *p* = 0.05, S2 Table), center distance (*F*_(1,47)_ = 9.4; *p* = 0.004, Fig 4E), and center time (*F*_(1,47)_ = 12.5; *p* = 0.001, Fig 4F). *Kif3a*^+/-^ genotype also had significant effects upon repeated beam breaks (*F*_(1,47)_ = 5.2; *p* = 0.03, S2 Table), center distance (*F*_(1,47)_ = 10.6; *p* = 0.002, Fig 4E), and center time (*F*_(1,47)_ = 7.4; *p* = 0.009, Fig 4F) Both NAE and *Kif3a*^+/-^ genotype significantly affected the number of rears, rotations, and total distance (statistics shown in S2 Table). On each of these measures, NAE *Kif3a*^+/-^ mice were the most active treatment group indicating an additive effect of *Kif3a* genotype and NAE.

To further illustrate the heightened activity of the NAE *Kif3a*^+/-^ mice relative to the other groups, as predicted by our hypothesis, we analyzed overall horizontal activity (Fig 4D), center distance (Fig 4E), and center time (Fig 4F) as 60-min totals and used Bonferroni *post hoc* tests to compare NAE *Kif3a*^+/-^ to NAE *Kif3a*^+/+^ mice (effect of genotype within NAE mice) or vehicle *Kif3a^+/-^* mice (effect of NAE within the heterozygotes), vehicle *Kif3a*^+/-^ to vehicle *Kif3a*^+/+^ (effect of genotype alone), and NAE *Kif3*a^+/+^ to vehicle *Kif3a*^+/+^ mice (effect of NAE within the wild-types). The NAE *Kif3a*^+/-^ mice were more active than their NAE *Kif3a*^+/+^ littermates (*p* = 0.001 for horizontal activity, 0.002 for center distance, and 0.006 for center time), and the vehicle *Kif3a*^+/-^ mice (*p* = 0.003 for horizontal activity, 0.015 for center distance, and 0.003 for center time). There were no significant effects of NAE or *Kif3a*^+/-^ alone. The heightened overall activity levels, as well as activity specific to the center of the open field which may reflect altered anxiety-like behavior, confirm our prior results from early NAE that open field activity is a sensitive index for the effects of early gestational alcohol exposure (11, 12). The greater sensitivity of the *Kif3a*^+/-^ in the open field versus the EPM is likely due to differences in the test duration because the differences between treatment and genotype groups appear to become greater as exploration patterns adapt to the environment.

Brain width and ventricle area were also analyzed in a subset of adolescent male mice (n = 4-5 per group) chosen based on performance closest to the group means on open field horizontal activity. Neither genotype nor prenatal treatment significantly affected ventricle area, though *Kif3a*^+/-^ mice tended to have larger ventricles (*F*_(1,13)_ = 3.767, *p* = 0.0743; Fig S5A). A genotype-treatment interaction was found for midline brain width (*F*_(1,13)_ = 9.206, *p* = 0.0096; Fig S5B), as well as a main effect of treatment (*F*_(1,13)_ = 27.55, *p* = 0.0002). Dunnett’s multiple comparison test was used to compare each group to vehicle-treated *Kif3*a^+/+^ animals, and it was determined that the brains in all three other groups were narrower (vs. *Kif3a*^+/+^ NAE: p = 0.0001; vs. *Kif3a*^+/-^ Veh: p = 0.0308; vs. *Kif3a*^+/-^ NAE: *p* = 0.0018). In addition, a main effect of genotype was found for medial brain height, with *Kif3a*^+/-^ mice having shorter medial height compared to *Kif3a*^+/+^ mice (*F*_(1,13)_ = 6.332, *p* = 0.0258; Fig S5C). Thus, the partial deletion of *Kif3a* resulted in smaller brains in adolescence that coincided with behavioral abnormalities.

## Discussion

The studies described here demonstrate that NAE disrupts the Shh pathway and cell cycle gene expression, and causes a reduction in RVNT volume during the first 24 hr following exposure. In addition, NAE disrupts expression of genes related to ciliogenesis and protein trafficking (Fig 4). Finally, cilia gene-alcohol interactions were further demonstrated through the use of *Kif3a* transgenic mice, a model of genetic ciliopathies (46). NAE male *Kif3a*^+/-^ exhibited exacerbated behavioral impairments on the open field and EPM compared to controls and WT NAE mice, implicating cilia in specific behavioral deficits. The current results support that early gestational alcohol exposure disrupts the Shh pathway, either independently or as a consequence of primary cilia dysregulation. Future work will focus on the exact impact of prenatal alcohol on primary cilia function and whether cilia dysfunction in the RVNT causes the observed downregulation of the Shh pathway and if impaired cilia function contributes to the development of craniofacial and midline CNS defects following prenatal alcohol exposure (11–14).

Additionally, these data have implications beyond the embryonic period and the neural tube, including informing the emerging study of enhanced cancer risk in human alcoholics. Dysregulation of the developmental signaling pathways, such as Shh, contribute to cancer pathogenesis (48) and more research is focusing on primary cilia dysfunction as a mediating factor in cancer development (49).

### NAE disrupts the Shh pathway and cilia function in the neural tube

These studies confirm that NAE downregulated Shh pathway expression, similar to other FASD models targeting early points of gestation (6–10). We observed decreased *Shh* and *Gli1* expression in the RVNT of NAE embryos during the first 24 hr post-exposure. These reductions coincided with increased levels of the Gli3^Rep^ form 12 hr following NAE. These data are the first to report altered Shh signaling following alcohol exposure during neurulation. While *Gli2* gene expression was not affected by NAE at either time point, it is possible that the ratio of the different forms of Gli2 was altered by NAE while not impacting levels of mRNA, similar to what was found for Gli3. Like Gli3, Gli2 is found in full-length activator and cleaved repressor forms, though studies have suggested that Gli2 acts primarily as a transcriptional activator in the mouse neural tube (50–52), meaning the majority of Gli2 will be found in the full-length form. However, currently available antibodies cannot reliably identify the various forms of Gli2, in part due to the dearth of antibodies that bind to the correct epitope of the protein to allow for detection of the cleaved repressor (N-terminus). It will be possible to address this question once validated antibodies targeting different amino acid sequences of mouse Gli2 become available.

The molecular mechanisms of action of alcohol on the Shh pathway remain unclear. Alcohol could act directly on the Shh pathway through interference with upstream regulator proteins, such as Hoxd’s (53), Sox2/3 (54), or Hand2 (best described in the embryonic limb bud) (55), or the Shh co-receptor Cdon (56), disruption of cholesterol (57) or other proteins necessary for Shh modulation, or activation of pathway inhibitors such as PKA (8) or Tbx2 (58). Our data suggest that alcohol could indirectly disrupt the Shh pathway as a downstream consequence of the observed dysregulation of primary cilia genes and their related functions. Downregulation of Gli proteins can cause a negative feedback loop in Shh expression as the Gli proteins target multiple members of the Shh path, including Ptc (59). The possibility that alcohol initiates a negative feedback loop of the Shh pathway is supported by the concomitant reduction in expression of pro-proliferative genes known to be downstream of Gli’s, *Ccnd1*, *Ccnd2*, and *Fgf15*. Expression of other targets of Gli-mediated transcription may have been impacted as well. The unique expression profiles of these genes during the 24 hr post-NAE indicate possible differences between these molecules in alcohol sensitivity or presence of compensatory mechanisms in the neural tube. The impact of reduced pro-proliferative genes was observed as a smaller RVNT volume at E9.5. Since previous work from our laboratory has shown little to no excessive apoptosis in the rostral basal or floor plates of the neural tube following E9.0 alcohol exposure (5), the most likely explanation for the reduced RVNT volume is decreased cell proliferation. The lack of increased apoptosis in the rostroventral portion of the neural tube following NAE stands in stark contrast to the pronounced apoptosis throughout the neuroectoderm following exposure at earlier time points and suggests region- and timing-specific mechanisms of alcohol-induced tissue damage. While not quantitatively measured in this study, the RVNT’s of NAE embryos displayed some shape differences compared to controls, as can be seen in Fig 2. Since neural patterning is a primary function of Shh, altered cell migration in the RVNT could be another consequence of disrupted Shh signaling. It is also possible that reduced activation of the Shh pathway in the ventral neural tube allows for redistribution of morphogens and increased expression of Wnt within the dorsal neural tube. Since the dorsal neural tube also contributes to craniofacial development through Wnt and Bmp signaling, an imbalance in the morphogenic gradients would likely result in abnormal growth trajectories of regions arising from these portions of the neural tube.

The Shh pathway requires properly functioning primary cilia and our data demonstrating NAE-induced disruptions to cilia gene expression support that NAE impairs primary cilia function. While cilia density was not affected in the current study, NAE resulted in changes to cilia-related genes in the RVNT that emerge within the first 6 hr following alcohol exposure and extend to the Gli family of transcription factors after 6-12 hr. These data demonstrate that NAE rapidly dysregulates primary cilia gene expression and identifies ciliogenesis, protein trafficking, and maintenance of cilia stability as targets of NAE. At E9.25, NAE upregulated *Hap1* and *Rilpl2* in the RVNT (Table 2; Fig 3C), two genes associated with ciliogenesis (60, 61). *Hap1* is named for its role in Huntington’s disease (60) and has been shown to mediate ciliogenesis through interaction with the Huntingtin protein (Htt). While little is known regarding ciliopathies in patients with Huntington’s disease, patients with Huntington’s, and mice with introduction of mutated Htt, exhibit abnormally increased cilia number and length (60, 62). Furthermore, mutated Htt disrupts the normal binding of Hap1 to dynein, affecting protein transport (63). In addition to its role in ciliogenesis, *Rilpl2* is involved in ciliary protein transport (61). Increased expression of ciliogenesis-related genes could be a result of alcohol-induced disruptions of normal cell cycle progression in the RVNT, as cilia are retracted and extended during mitosis.

In contrast, *Cep41*, *Nek4*, *Evc*, and *Dpcd* were downregulated in NAE embryos at E9.25 (Table 2, Fig 3C). *Cep41* is known to be mutated in patients with the genetic ciliopathy Joubert syndrome (64). Patients with Joubert syndrome display many symptoms similar to other ciliopathies, including renal, ocular, digital, and neurological abnormalities. Notably, patients with Joubert syndrome also show clinical symptoms that overlap with some features of FASD, including cerebellar dysplasia, impaired motor function, abnormal eye development, polydactyly, and craniofacial dysmorphologies. In mouse and zebrafish models, loss of *Cep41* causes ciliopathy-like phenotypes, including brain malformations (64). Thus, even transient downregulation of this gene could impact normal CNS development. In addition, Cep41 is implicated in polyglutamylation of ciliary α-tubulin; loss of tubulin glutamylation could result in structural instability and affect ciliary assembly and transport. Further studies are needed to determine if changes to *Cep41* expression are directly linked to glutamylation status of ciliary tubulin in the RVNT following NAE. *Nek4* has a role in mediating ciliary assembly and integrity, possibly through *Nek4*’s regulation of microtubules (65). The gene *Dpcd* is associated with another ciliopathy, Primary Ciliary Dyskinesia, caused by dysfunction of motile cilia in the respiratory tract and reproductive systems, resulting in respiratory difficulties and infertility (66). *Dpcd* has been implicated in the formation and maintenance of cilia and is upregulated during cell division. Finally, downregulation of *Evc* could have direct impact on trafficking of the Shh pathway proteins Smo and the Gli family (67, 68). Ellis-van Creveld syndrome, caused by a mutation in *Evc*, is a genetic ciliopathy presenting with polydactyly and other digital anomalies, congenital heart defects, and other skeletal abnormalities. Similar skeletal malformations have been noted in humans and rodent models of FASD (69–71). The decrease in *Cep41* was particularly interesting, as polyglutamylation of tubulin is important for protein trafficking within the cilia, including Shh pathway proteins (72). In addition, *Evc* is known to play a role in Gli protein trafficking (67). Together, the downregulation of these two genes suggests disruption of Gli signaling within the cilia as a likely mechanism for the later reduction in *Ccnd1* expression. The observed downregulation of *Dpcd,* a gene normally upregulated during cilia formation and cell division (66), further supports the conclusion that the cell cycle has been affected by NAE as early as 6 hr post-exposure. Transient changes in cell proliferation in this region of the neural tube would impact the development of midline brain and craniofacial tissue (4, 11, 13, 14). A schematic of cilia genes altered following NAE at 6 and 12 hr is shown in Fig 5.

**Fig 5.**
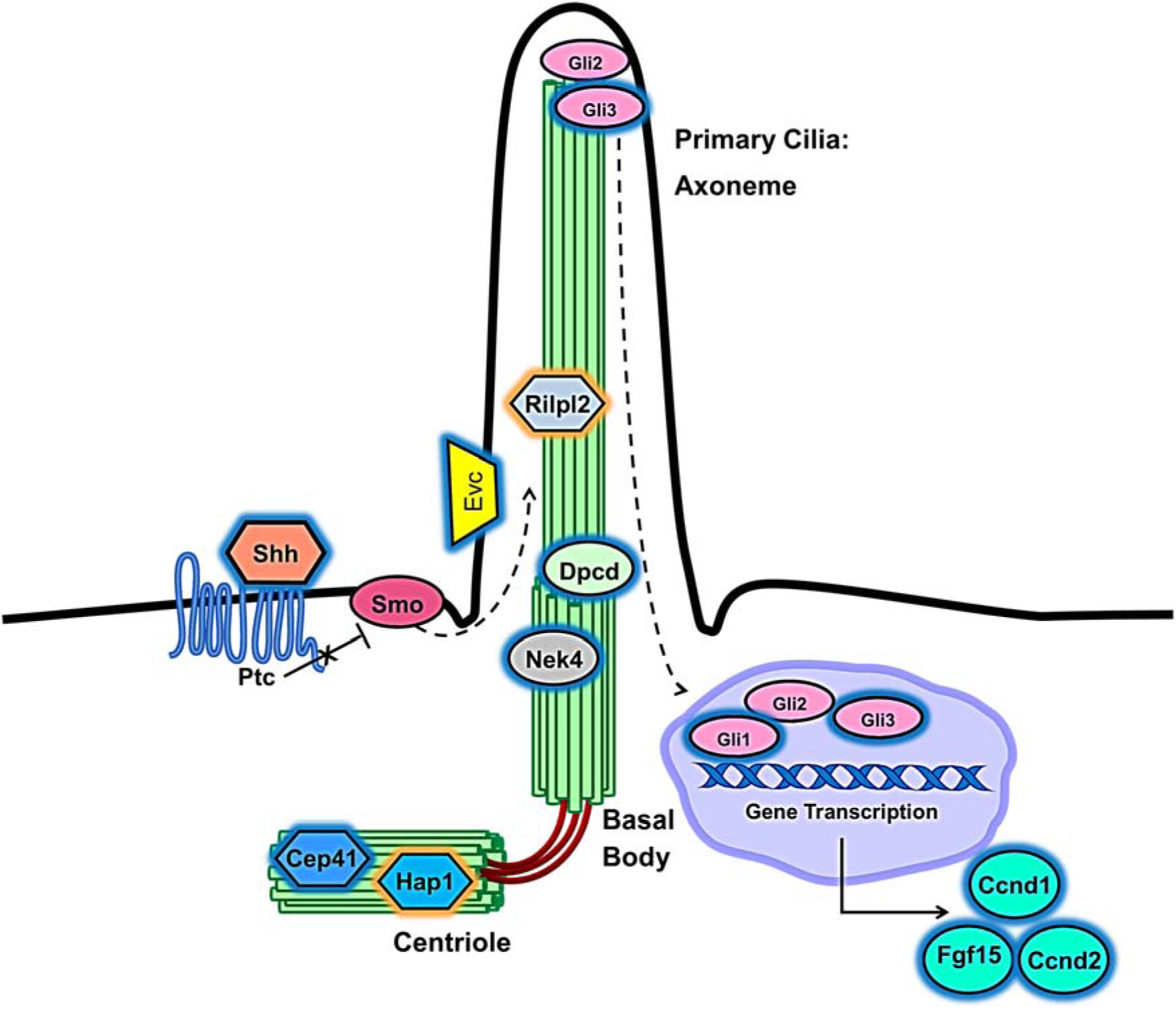
Schematic of a primary cilium. This representation of the primary cilium includes the axoneme, basal body, and centriole and displays differentially expressed genes related to cilia structure/ciliogenesis/protein trafficking 6 hr post-exposure and the Shh pathway and cilia function 6-24 hr post-exposure. Orange outlines indicate upregulated genes whereas blue outlines denote genes that were downregulated by NAE.

### NAE interacts with cilia gene knockdown to alter exploratory behavior during adolescence

Mice lacking one copy of the *Kif3a* gene were used to demonstrate that prenatal alcohol interacts with cilia function. The *Kif3a* mutant mouse is a well-characterized model of genetic ciliopathy (24, 46, 73) as *Kif3a* mutant mice phenocopy many human ciliopathies, including hypertelorism, facial clefting, and brain abnormalities (46, 74). Similar to the defects seen in conditional *Kif3a* mutant mice, NAE mice can exhibit abnormal cortical hemispheres and ventral midline brain structures (13, 14) and hypertelorism has been reported in some patients with heavy prenatal alcohol exposure who lack the classic FAS facial features (4, 75). *Kif3a* heterozygous mice have been shown to display similar physical abnormalities, such as situs inversus, to the full knockout but at a much lower rate (76), as *Kif3a*^-/-^ die by E10. *Kif3a*^+/-^ mice also have reported reduced *Kif3a* gene expression in the bone and osteoblasts at 6 weeks old (77). These mice also display abnormal *Gli2* expression (77), supporting that Kif3a interacts with Shh signaling and that *Kif3a*^+/-^ mice have abnormal cilia function, as Gli2 processing is a critical function of primary cilia during development. In the current study, exploratory behavior in two novel environments (EPM and open field), which relies heavily on the cortico-limbic and hypothalamic circuitry derived from the RVNT, was sensitive to both NAE and *Kif3a* genotype effects, as well as NAE-genotype interactions. NAE and *Kif3a*^+/-^ similarly heightened exploration of the EPM open arms and the center of the open field and increased total activity on the EPM and in the open field. Increased exploratory behavior suggests an impaired stress and or anxiety-related response, as previously observed following gestational alcohol exposure (35). While the current evidence cannot definitively show that the perturbed behaviors observed in NAE and *Kif3a*^+/-^ mice are due to a common mechanism, the persistence of hyperactivity and impaired open field habituation observed in the NAE *Kif3a*^+/-^ mice lends support to the hypothesis that variants of a key primary cilia gene might act as a risk factor for certain behavioral effects of NAE.

## Conclusions

In conclusion, these data demonstrate that neurulation-stage alcohol exposure decreases Shh pathway signaling, Shh-mediated cell cycle gene expression, and alters expression of genes related to ciliogenesis and protein trafficking in a region of the neural tube that gives rise to alcohol-sensitive ventral midline brain structures. These results are the first to suggest primary cilia dysfunction as a possible pathogenic mechanism of prenatal alcohol exposure, either through downstream effects on the Shh pathway or other mechanisms that remain to be elucidated. Furthermore, NAE interacts with primary cilia to alter open field and EPM behavior in *Kif3a* mutant male mice during adolescence. Together, these data demonstrate an interaction of prenatal alcohol and primary cilia, possibly resulting in an alcohol-induced “transient ciliopathy” and contributing to midline craniofacial and CNS defects in FASD.

## Acknowledgements

We thank Jamie Leitzinger, Divya Venkatasubramanian, Haley Mendoza-Romero, Laura Murdaugh, and Debbie Dehart for their technical assistance on this project.

## Competing interests

The authors have no conflicts of interest to report.

## Funding

Funding to support this research was provided by the National Institutes of Health/National Institute of Alcohol and Alcoholism [U01AA021651 and R01AA026068 to SEP, F32AA026479 to KEB] and conducted as part of the Collaborative Initiative on Fetal Alcohol Spectrum Disorders (CIFASD).

## Author Contributions

Conceptualization: KEB, EWF, and SEP

Methodology: KEB, EWF, and SEP

Investigation: KEB and EWF

Formal Analysis: KEB and EWF

Visualization: KEB and EWF

Writing – Original Draft Preparation: KEB and EWF

Writing – Review & Editing: KEB, EWF, and SEP

## Supporting information

**S1 Fig. Uncropped western blot analysis of Gli3**. A) E9.25 (6 hr post-exposure) blot of Gli3 full length (190 kDa), cleaved (83 kDA), and Gapdh (37 kDa) bands. B) E9.5 and C) E10 blot of Gli3 and Gapdh. NAE: Neurulation-stage alcohol exposure; Veh: Lactated Ringer’s vehicle group. Each lane represents one pooled litter from the treatment group indicated.

**S2 Fig. Representative images of NAE (A) and vehicle-treated (B) embryos 12 hr after exposure (E9.5).** No gross morphological differences were observed between the treatment groups at this time point, though volumetric and shape differences of CNS tissue are apparent later in development (11–14). Scale bars = 1 mm.

**S3 Fig. NAE causes impairments in rotarod performance in male, but not female, mice.** Male NAE mice, regardless of genotype, had a significantly shorter latency to fall compared to vehicle-treated mice (*F*_(1,48)_ = 5.1; *p* = 0.028). No effect of genotype or treatment on rotarod performance was found in female mice. A main effect of trial was found (*F*_(4,220)_ = 24.46, *p* < 0.0001), demonstrating motor learning in the females across trials. * = *p* < 0.05. For males, n’s = 11-15 litters; for females, n’s = 13-17 litters. Data are shown as group means ± SEM with each sample representing littermates of each genotype averaged into a single datum.

**S4 Fig. Partial loss of cilia motor transport gene *Kif3a* affects behavioral performance on the EPM (A-C) and open field (D-F) in adolescent female mice.** For EPM, total arm entries (A) and percent of time (B) in open arms showed no effects of treatment or genotype. However, C) a significant main effect of genotype was revealed for percent of entries into open arms (*F*_(1, 57)_ = 5.13, *p* = 0.028), with *Kif3a*^+/-^ female mice making more entries into open arms compared to WT mice. For open field, there was a significant genotype effect (indicated by curved brackets) on horizontal activity (D) (*F*_(1,54)_ = 9.2, *p* =0.004), center distance (E) (*F*_(1,54)_ = 7.9, *p* = 0.007), center time (F) (*F*_(1,54)_ = 6.4; *p* = 0.014), and repeated beam breaks (S1 Table) (*F*_(1,54)_ = 6.7; *p* = 0.013). Overall, when all time bins were averaged, *Kif3a*^+/-^ female mice were more active and traveled farther in the center of the open field compared to WT females. *Post hoc* analyses were not significant. * = *p* < 0.05, ** = *p* < 0.01. When data were totaled across the session (D-F), Bonferroni *post hoc* tests revealed that Veh *Kif3a*^+/-^ mice had more beam breaks (*p* = 0.005) and center distance traveled (*p* = 0.003) compared to Veh WT mice. For totaled data, significant (*p* < 0.05) *post hoc*s are shown as letters on each graph. a: vs. vehicle-treated WT, b: vs. vehicle-treated *Kif3a*^+/-^, c: vs. NAE WT. For all groups, n’s = 13-17 litters. All data are shown as group means ± SEM with each sample representing the females from one litter.

**S5 Fig. Partial loss of *Kif3a* reduces brain width and height in adolescent male mice.** A) Neither genotype nor prenatal treatment affected ventricle area. However, B) midline brain width was smaller in both *Kif3a*^+/-^ groups and NAE WT animals compared to vehicle-treated WT mice. C) *Kif3a* heterozygosity also results in smaller medial brain height. * = *p* < 0.05, ** = *p* < 0.01, *** = *p* < 0.001. Group n’s = 4-5. All measurements shown as mean + SEM.

**S1 Table. Open field behavioral outcomes in adolescent female *Kif3a*^+/-^ and *Kif3a*^+/+^ mice following NAE or vehicle treatment on E9.0.** ^a^ = Veh *Kif3a*^+/-^ and NAE *Kif3a*^+/-^ (averaged across time bins) significantly differ from the average of Veh WT.

**S2 Table. Open field behavioral outcomes in adolescent male *Kif3a*^+/-^ and *Kif3a*^+/+^ mice following NAE or vehicle treatment on E9.0.** ^a^ = Veh *Kif3a*^+/-^ and NAE *Kif3a*^+/-^ (averaged across time bins) significantly differ from the average of Veh WT.

