## Supplementary figures and images for "Prenatal alcohol exposure disrupts Shh pathway and primary cilia genes in the mouse neural tube"

### Supp Fig 1

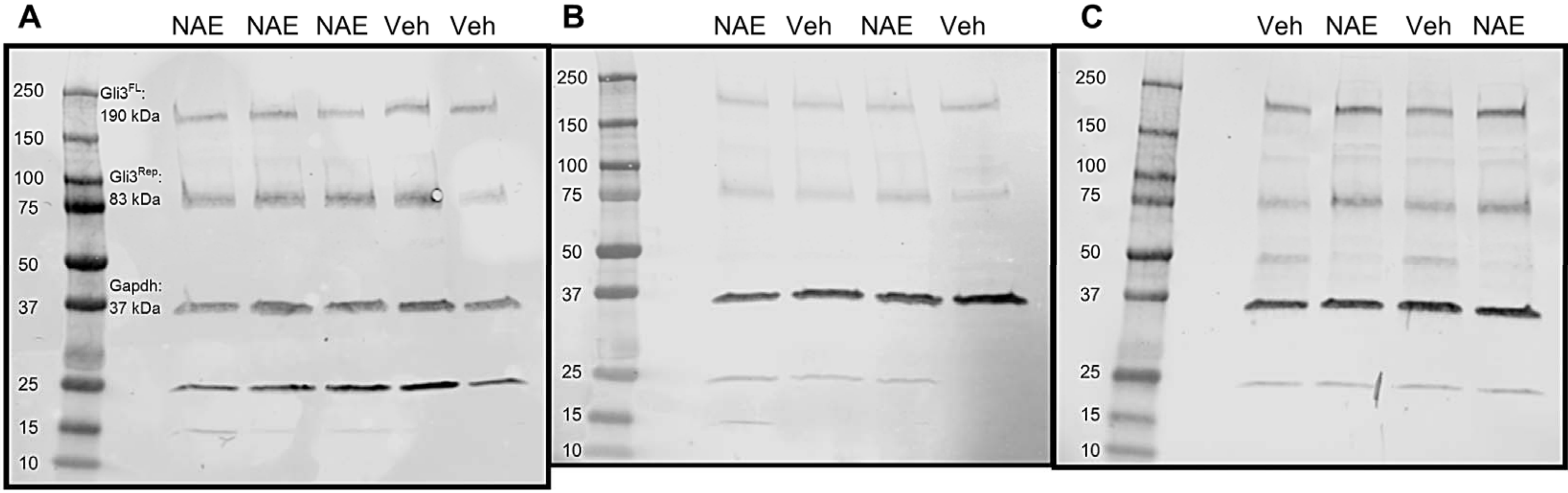

### Supp Fig 2

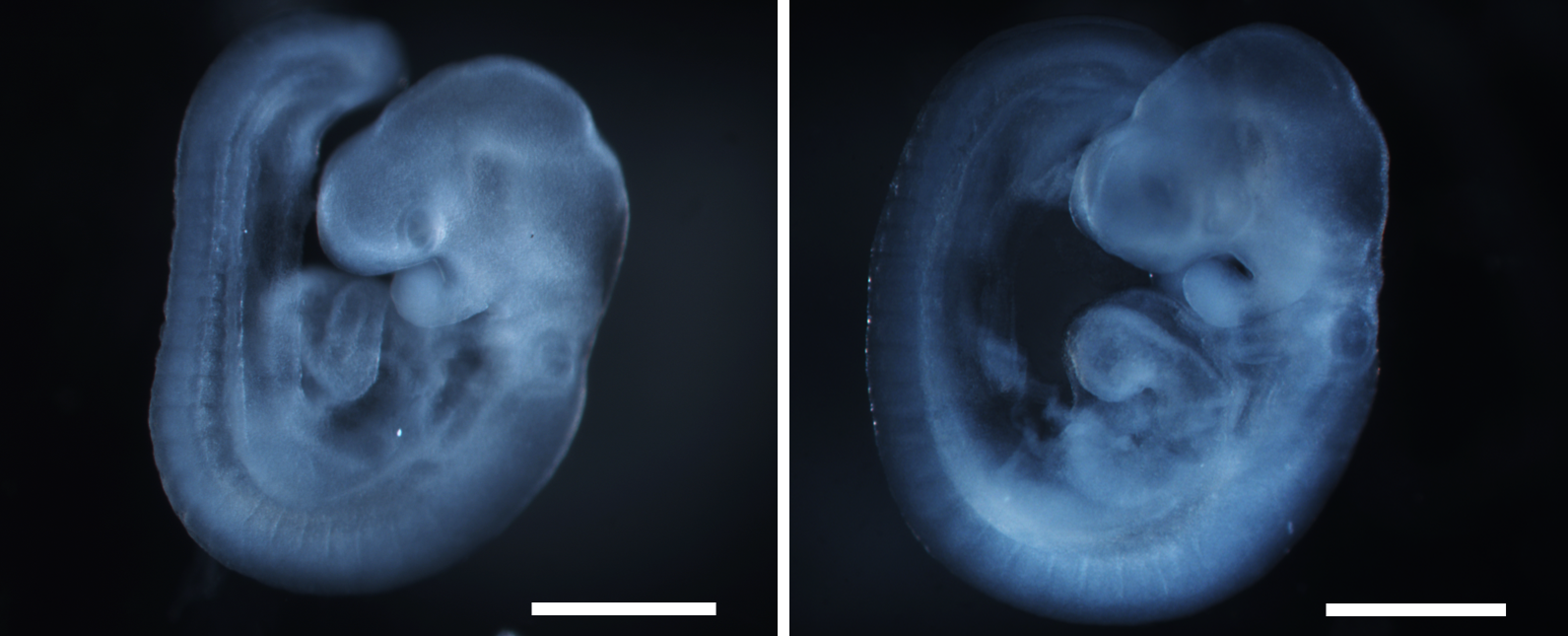

### Supp Fig 3

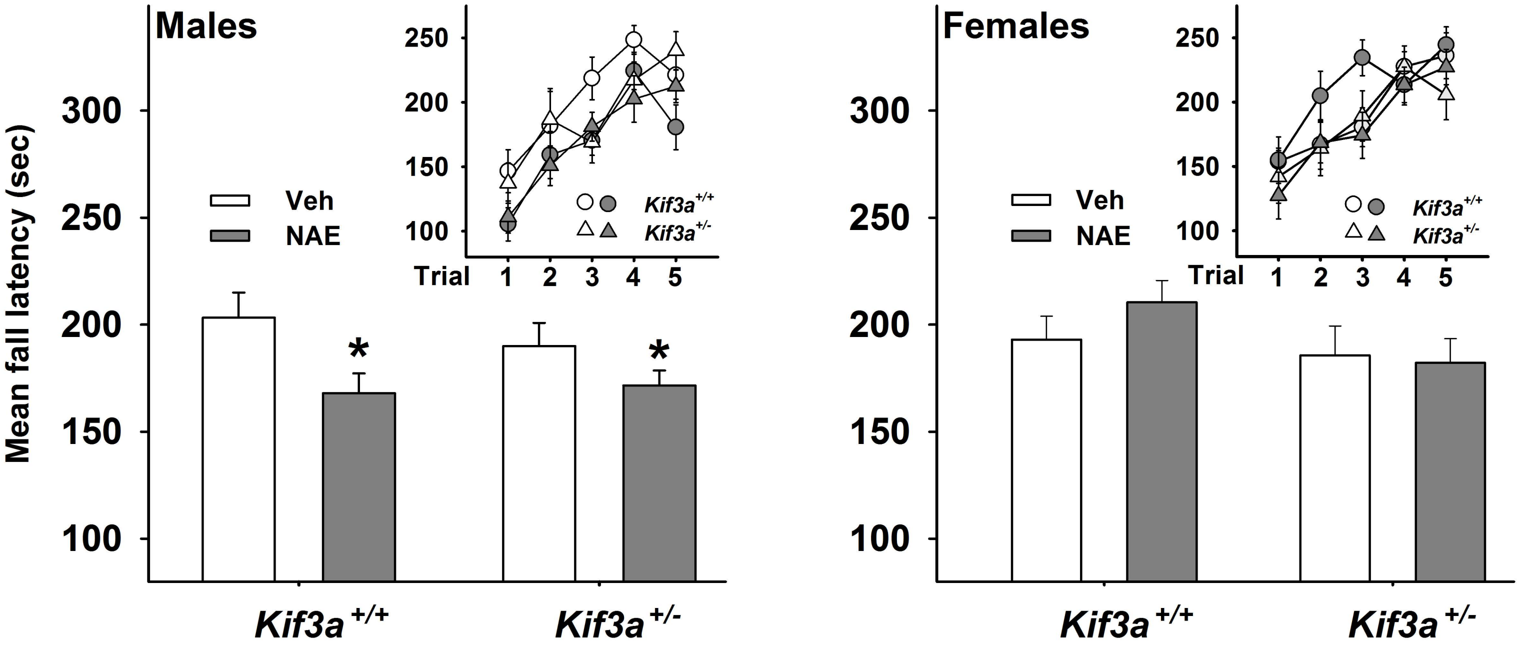

### Supp Fig 4

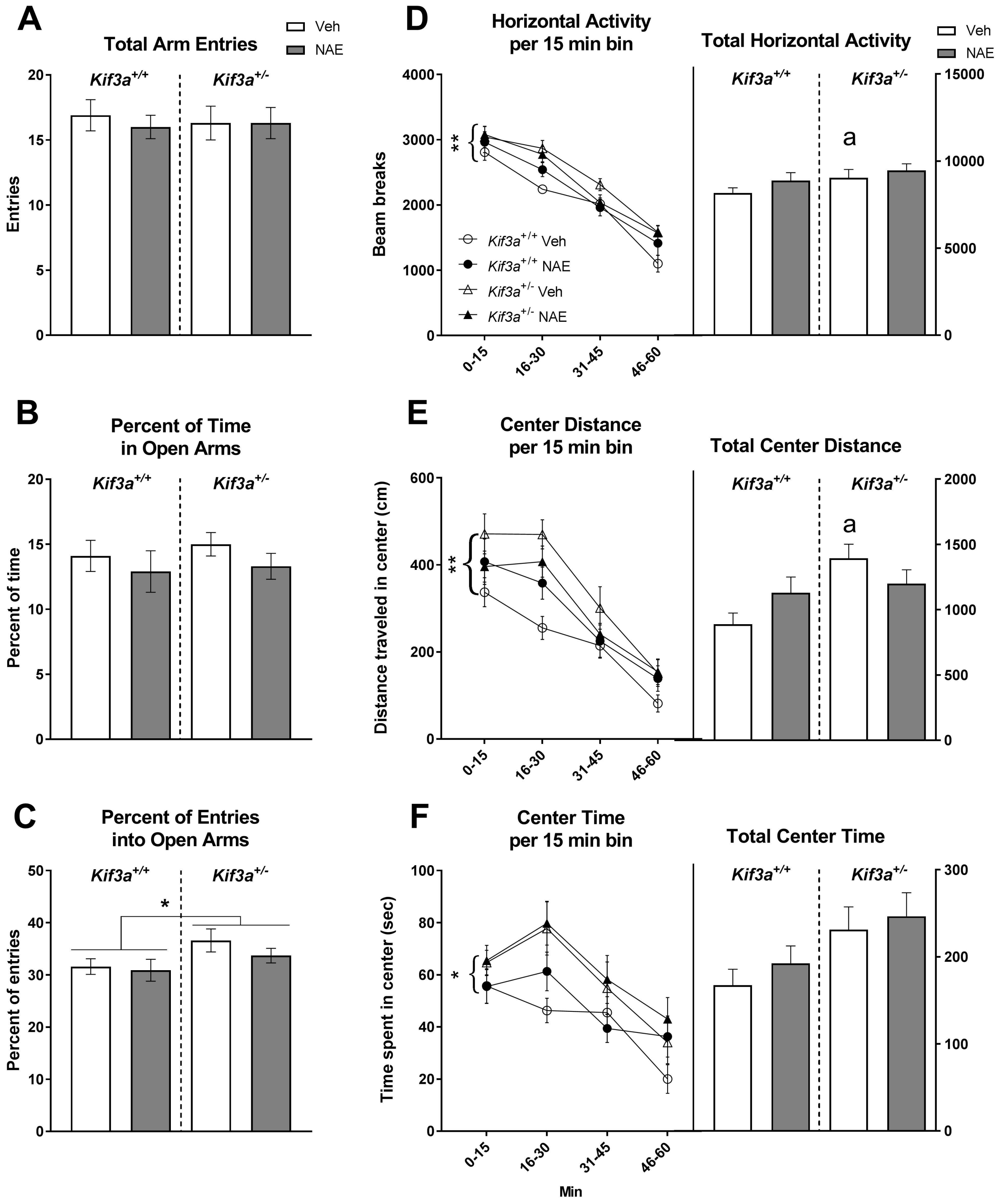

### Supp Fig 5

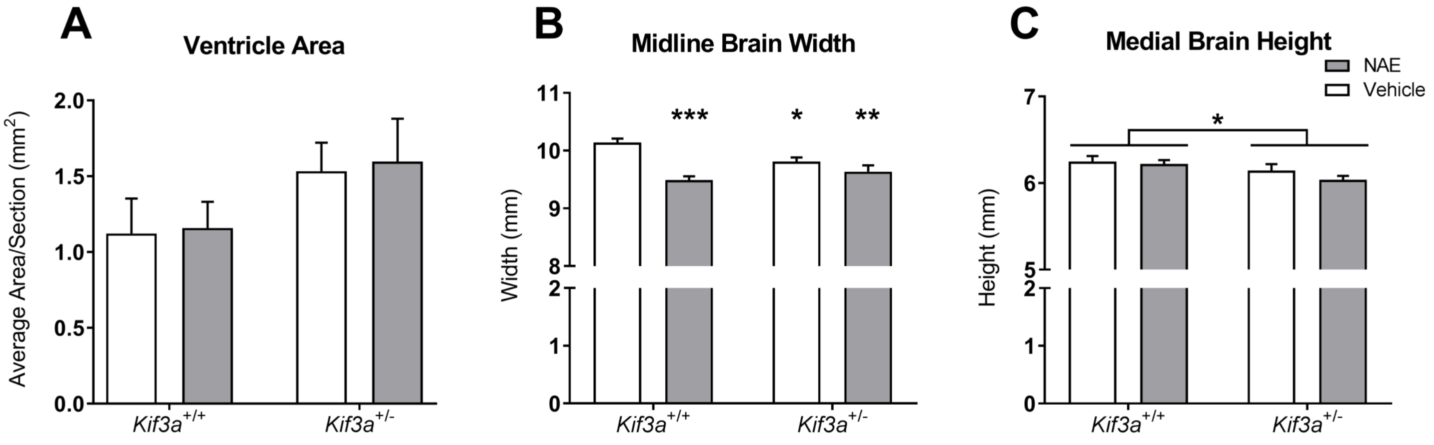
