## Supplementary material for "Prenatal alcohol exposure disrupts Shh pathway and primary cilia genes in the mouse neural tube": Supp Table 1

Supplemental Table 1. Open field behavioral outcomes in adolescent female *Kif3a*$^{+/-}$ and *Kif3a*$^{+/+}$ mice following NAE or vehicle treatment on GD9.0. [a] = Veh *Kif3a*$^{+/-}$ and NAE *Kif3a*$^{+/-}$ (averaged across time bins) significantly differ from the average of Veh WT.

| Measure | Effect | F *(df)* | *p*-value | Means ± SEM | | | | |
|---|---|---|---|---|---|---|---|---|
| | | | | **Groups** | **0-15 min** | **15-30 min** | **30-45 min** | **45-60 min** |
| **Number of rears** | Time x Treatment x Genotype Interaction | 6.1 (1, 54) | 0.001 | Veh WT: | 53.13±2.42 | 40.65±2.62 | 28.71±2.01 | 12.44±2.06 |
| | | | | NAE WT: | 57.66±3.29 | 48.0±3.75 | 29.57±2.49 | 18.9±3.26 |
| | | | | Veh *Kif3a*$^{+/-}$: | 43.38±4.06 | 50.79±3.92 | 35.93±2.71 | 21.95±3.21 |
| | | | | NAE *Kif3a*$^{+/-}$: | 52.08±3.35 | 49.4±2.59 | 31.58±2.8 | 17.92±2.14 |
| **Rotations** | Treatment x Genotype Interaction | 4.8 (1, 54) | 0.033 | Veh WT: | 12.44±0.77 | 8.45±0.47 | 5.73±0.54 | 3.21±0.64 |
| | | | | NAE WT: | 12.45±.095 | 10.22±0.71 | 6.27±0.71 | 4.43±0.74 |
| | | | | Veh *Kif3a*$^{+/-}$: | 15.36±1.23 | 12.23±0.92 | 8.56±0.57 | 5.60±0.61[a] |
| | | | | NAE *Kif3a*$^{+/-}$: | 13.98±0.60 | 10.53±0.79 | 7.0±0.77 | 4.52±0.61[a] |
| **Total distance** | Genotype | 7.9 (1, 54) | 0.007 | Veh WT: | 1809.26±88.28 | 1187.41±42.59 | 963.99±65.37 | 449.67±72.60 |
| | | | | NAE WT: | 1931.59±117.89 | 1427.52±89.83 | 1008.78±83.74 | 696.4±116.36 |
| | | | | Veh *Kif3a*$^{+/-}$: | 2100.76±135.51 | 1636.45±66.51 | 1228.51±85.15 | 761.02±73.75 |
| | | | | NAE *Kif3a*$^{+/-}$: | 1955.03±66.97 | 1599.32±82.39 | 1039.10±97.35 | 712.77±92.86 |

Supplemental Table 2. Open field behavioral outcomes in adolescent male *Kif3a*$^{+/-}$ and *Kif3a*$^{+/+}$ mice following NAE or vehicle treatment on GD9.0.

| Measure | Main effect | F *(df)* | *p*-value | Means ± SEM | | | | |
|---|---|---|---|---|---|---|---|---|
| | | | | **Groups** | **0-15 min** | **15-30 min** | **30-45 min** | **45-60 min** |
| **Number of rears** | Treatment | 12.1 *(1, 47)* | 0.001 | Veh WT: | 41.8±2.99 | 46.35±3.52 | 35.62±5.68 | 20.94±4.56 |
| | | | | NAE WT: | 53.81±2.73 | 54.72±3.03 | 47.75±3.37 | 23.59±4.33 |
| | Genotype | 6.8 *(1, 47)* | 0.012 | Veh *Kif3a*$^{+/-}$: | 48.17±3.62 | 38.55±4.23 | 38.55±4.23 | 27.99±5.48 |
| | | | | NAE *Kif3a*$^{+/-}$: | 58.94±4.19 | 46.26±4.24 | 46.26±4.23 | 41.55±3.37 |
| **Rotations** | Treatment | 6.9 *(1, 47)* | 0.01 | Veh WT: | 10.33±0.57 | 8.0±0.62 | 5.25±0.62 | 4.24±0.88 |
| | | | | NAE WT: | 12.30±0.75 | 8.74±0.45 | 8.33±0.74 | 4.96±0.94 |
| | Genotype | 8.2 *(1, 47)* | 0.006 | Veh *Kif3a*$^{+/-}$: | 11.95±0.93 | 11.17±0.97 | 6.68±0.77 | 5.08±0.86 |
| | | | | NAE *Kif3a*$^{+/-}$: | 13.36±0.52 | 11.12±0.64 | 9.09±1.16 | 6.79±0.77 |
| **Total distance** | Treatment | 7.0 *(1, 47)* | 0.01 | Veh WT: | 1522.88±70.66 | 1274.44±46.22 | 956.55±114.24 | 631.62±120.82 |
| | | | | NAE WT: | 1816.743±89.16 | 1371.74±48.36 | 1104.47±110.44 | 695.46±132.27 |
| | Genotype | 5.0 *(1, 47)* | 0.03 | Veh *Kif3a*$^{+/-}$: | 1687.403±75.98 | 1463.0±74.32 | 1010.12±86.27 | 678.29±105.14 |
| | | | | NAE *Kif3a*$^{+/-}$: | 1989.38±100.53 | 1632.72±116.49 | 1280.71±134.02 | 1096.01±152.89 |
